# B cell expression of the enzyme PexRAP, an intermediary in ether lipid biosynthesis, promotes antibody responses and germinal center size

**DOI:** 10.1101/2024.10.17.618760

**Authors:** Sung Hoon Cho, Marissa A. Jones, Kaylor Meyer, David M. Anderson, Sergei Chetyrkin, M. Wade Calcutt, Richard M. Caprioli, Clay F. Semenkovich, Mark R. Boothby

## Abstract

The qualities of antibody (Ab) responses provided by B lymphocytes and their plasma cell (PC) descendants are crucial facets of responses to vaccines and microbes. Metabolic processes and products regulate aspects of B cell proliferation and differentiation into germinal center (GC) and PC states as well as Ab diversification. However, there is little information about lymphoid cell-intrinsic functions of enzymes that mediate ether lipid biosynthesis, including a major class of membrane phospholipids. Imaging mass spectrometry (IMS) results had indicated that concentrations of a number of these phospholipids were substantially enhanced in GC compared to the background average in spleens. However, it was not clear if biosynthesis in B cells was a basis for this finding, or whether such cell-intrinsic biosynthesis contributes to B cell physiology or Ab responses. Ether lipid biosynthesis can involve the enzyme PexRAP, the product of the *Dhrs7b* gene. Using combinations of IMS and immunization experiments in mouse models with inducible *Dhrs7b* loss-of-function, we now show that B lineage-intrinsic expression of PexRAP promotes the magnitude and affinity maturation of a serological response. Moreover, the data revealed a *Dhrs7b*-dependent increase in ether phospholipids in primary follicles with a more prominent increase in GC. Mechanistically, PexRAP impacted B cell proliferation via enhanced survival associated with controlling levels of ROS and membrane peroxidation. These findings reveal a vital role of this peroxisomal enzyme in B cell homeostasis and the physiology of humoral immunity.

## INTRODUCTION

The qualities and concentrations of antigen (Ag)-specific antibodies (Ab) are key elements of immunity and the pathophysiology of diverse conditions involving inflammation (1–3). Ab are secreted by cells in the B lymphocyte lineage, but the exact sources are diverse (4–7). Many derive from a subset termed B1 B cells, which are thought to having characteristics of innate responses (4, 5), but progeny of follicular and marginal zone B cells (FoB and MZB, respectively) can differentiate into Ab-secreting plasma cells (PC) after activation and proliferation (6–8). Ab concentrations and affinity for Ag, as well as Ab isotype after class switching in a B cell, determine the efficacy of pathogen clearance (9–12). Importantly, the affinities of serum Ab for target Ag can increase over time after an immune exposure, especially after recurrent encounter(s) (8, 9, 13, 14).

Many sources and forms of Ab can be protective, though in several autoimmune diseases these molecules and the somatic mutations that diversify an initial BCR repertoire drive pathology (15, 16). The micro-anatomic structure termed the germinal center (GC) provides one mechanism to increase spectrum of Ab affinities over time and repeated Ag exposure (17–19). Although T cell help-dependent diversification and PC production can occur independent from GC, results with different forms of vaccination support the practical importance of GC (20, 21). Current evidence holds that most Ig class-switching is executed before GC entry of proliferating B cells (22). GC reactions are initiated after B cells in a primary lymphoid follicle encounter Ag, migrate to interact with activated helper T cells after extensive proliferation (23). GC formation, size, and function depend on multiple factors. These include the efficiency of population growth for B cells in a phase before they enter and take on characteristics of GC B cells, and on homeostatic and differentiative processes while in the secondary follicle [reviewed in (24)]. Once in the GC, B cells cycle between light and dark zones, undergoing iterative cycles of selection. Proliferation and selection end in development as Ab-secreting plasma cells or memory B cells [reviewed in (14, 18, 19)]. In line with a selection process, substantial rates of B cell death have been measured in GC (25). As such, the qualities and quantities of Ab elicited by immunization (or vaccination) depend on B cell proliferation and survival both before and during GC reactions.

Accumulating evidence indicates that metabolic reprogramming and intermediary metabolism play pivotal roles in survival, proliferation, differentiation and function in the pre-immune populations (26, 27), including B cells (28, 29). Among core metabolic processes, several converge on oxidative metabolism fed by glycolytic generation of pyruvate, anaplerotic use of amino acids such as glutamine, and fatty acid oxidation [reviewed in (29, 30)]. Compared to resting populations such as naive B cells, rates of such metabolism in B cells appear to be increased after activation, including in GC B cells [31-36; reviewed in (29)]. Such increases support the generation of substrates for anabolic processes in the dividing cell population and perhaps the energy demands thereof.

Mitochondria have been the focus of investigations exploring molecular mechanisms for oxidative metabolism to generate energy and substrates in lymphocytes during immune responses [reviewed in (37, 38)]. Oxidative phosphorylation in and dynamics of this organelle appear to be important in determining rates of somatic hypermutation that characterize GC B cell biology (33–35), albeit by a molecular mechanism that has not yet been identified. Among specialized functions of this highly dynamic subcellular organelle, mitochondria use the Krebs TCA cycle to feed an electron transport chain (ETC) and efficiently generate ATP. Along with other sources, mitochondria generate reactive oxygen species (ROS) via the ETC at a rate determined by structural and biochemical features (39, 40). Rates of ROS production, release, and resolution require regulation at multiple levels because these labile species and their reactions with cellular molecules are important for both signaling and cell proliferation but undermine cell function or survival when excessive.

B cells in most GC exhibit inadequacy of oxygen delivery relative to demands (31, 41–43). Of note, genome-wide screening for genes that help cells adapt to such hypoxia revealed that peroxisomes and an ER enzyme involved in synthesizing plasmalogen ether lipids (EL) decrease the death of cells due to insufficient oxygen (44). Using an unbiased lipidomic analysis that combined imaging mass spectrometry (IMS) and techniques for identifying GC in sections of spleen from immunized mice, we had found that local concentrations of a subset of EL were heightened in GC relative to other portions of the spleen (45). Ether phospholipids (EPL) have a fatty acid linked to the glycerol backbone with an alkyl or a vinyl ether bond at the sn-1 position (46, 47). Synthesis of ether phospholipids depends on the generation of precursors in peroxisomes and final processing in the ER (48, 49). However, EL biosynthesis is prominent in the liver, and in theory these species or their precursors might distribute to tissues from a primary site of biosynthesis or from the diet (49–53). Thus, the finding that secondary follicles had greater densities of some - but not all - ether phospholipids left several key questions unanswered – (1) Does the capacity to synthesize and increase ether lipids in B cell follicles – primary or secondary (i.e., GC) - affect antibody responses? (2) Are any of the EL dependent on B lineage-intrinsic metabolism to generate precursors or final products? Moreover, there is debate whether EL or their plasmalogen subset promote cell death or resistance, and whether inactivation of biosynthesis causes defects in establishment or maintenance of hematopoietic cells (46, 53–55).

Peroxisomal Reductase Activating PPARγ (PexRAP, encoded by the *Dhrs7b* gene) catalyzes the reduction of alkyl-dihydroxyacetonephosphate (DHAP) to 1-alkyl-G3P (46, 48, 56), which then is transferred into endoplasmic reticulum (ER) and further metabolized to generate EL. Recent work notes that this enzyme may substantially function in the ER in addition to the peroxisome (56). Widespread inactivation of *Dhrs7b* in mature mice decreased hepatic production of ether lipid species and reduced the population of erythrocytes and neutrophils (46). This body of work indicated that PexRAP can be essential in some aspects of ether lipid metabolism and may function in hematopoietic cells, but was questioned based on inborn errors of metabolism affecting other aspects of ether lipid biosynthesis (54). Diverse functions of ether lipids in general, and the plasmalogen subset in particular, have been proposed or supported by prior work [reviewed in (57–61)]. The ether linkage may provide a sink for oxygen radicals - thereby acting directly to resolve the reactivity and contribute to ROS homeostasis, although there is debate as to the physiological impact of this chemical property (62). Whether or not this lipid subset influences adaptive immunity or conventional lymphocyte lineages, however, is unclear.

Herein, we tested the hypotheses that (i) the ether lipid profiles in activated B cells depend on their expression of the biosynthetic enzyme PexRAP [alternatively referred to as acyl/alkyl-DHAP reductase (ADHAPR) (56)], and (ii) this enzyme promotes B cell physiology. To do so, we combined conventional lipidomics, Imaging Mass Spectrometry (IMS), and immunization experiments with genetic models in which loss-of-function for *Dhrs7b* is induced in adult mice. We provide evidence that PexRAP impacts B cell proliferation via enhanced survival associated with controlling levels of ROS and membrane peroxidation. The results also indicate that beyond an impact on B cell homeostasis, GC are reduced in its absence, and the magnitude and affinity maturation of a serological response depend on B cell expression of PexRAP. Taken together, these findings support a function of B cell-intrinsic biosynthesis of ether lipids in B lymphocyte homeostasis, the production of Ab, and in promoting GC reactions.

## RESULTS

### PexRAP promotes homeostatic maintenance and proliferation of B cells

By use of IMS, we had reported that at least a dozen ether phospholipids - including plasmalogens - were enriched in splenic germinal centers of immunized mice when compared to the rest of the tissue (45). Previous work provided evidence that these lipid molecules were important for the homeostasis of short-lived neutrophils (46). However, this effect was questioned based on hematological data from patients inborn errors of metabolism that affect ether lipid biosynthesis (54). Moreover, the impact of these molecules on adaptive immune cells or functions is not known. Accordingly, we sought to test for effects of an intervention that interferes with a key biosynthetic pathway used for endogenous synthesis of these molecules. To do so, we started with analyses using induced loss-of-function for *Dhrs7b^f/f^* and the widely expressed *Rosa26-CreER^T2^* transgene (63, 64) used in (46), which reduced hepatic generation of a subset of ether lipids and acts in all hematopoietic cells as well.

We modeled initial experiments on the published work with imaging mass spectrometry (45). Mice were treated with tamoxifen using conditions less intense than a regimen shown to have no effect on pre-immune splenic and B lineage populations (65, 66), immunized with SRBC [akin to the analyses in (31, 36, 41, 45)], and analyzed 7 d after immunization (Fig. 1A). Functional inactivation of *Dhrs7b* in T and B lymphocytes was confirmed by Western blot analysis using anti-PexRAP Ab (Fig. 1B). The frequencies and numbers of CD19^+^ B220^+^ B cells in *Dhrs7b^f/f^*; *Rosa26-CreER^T2^* - only a small minority of which would have been activated by the immunization - were approximately 0.6 the values for controls (*Dhrs7b*^+/+^ *Rosa26-CreER^T2^*, i.e., wild-type locus) (Fig. 1C, D). In contrast to B cell populations, TCRβ^+^ CD4^+^ T cells and TCRβ^+^ CD4^-^CD8 T cells were intact in tamoxifen-treated and SRBC-immunized *Dhrs7b^f/f^*; *Rosa26-CreER^T2^* mice (Supplemental Fig. 1A-C). These results suggest that *Dhrs7b* is required for fully maintaining the B cell population but indicate that the enzyme is dispensable for the main sets of conventional T cells. In light of the earlier finding that a subset of ether lipids was more prevalent in secondary follicles, we analyzed GC in these samples. Microscopy and flow cytometry analyses after immunofluorescent staining found that less GC B cells were present after immunization of *Dhrs7b^f/f^*; *Rosa26-CreER^T2^* mice, i.e., less IgD^-^CD95^+^ GL7^+^ (GC-phenotype) B cells (Fig. 1E-I; Supplemental Fig. 1D). Quantitation of the immune fluorescence micrography with spleens from immunized mice revealed that the numbers of GC per spleen were halved in *Dhrs7b*^Δ/Δ^ mice compared with controls (Fig. 1I; Supplemental Fig. 1E). The sizes of those GC that did form were substantially reduced as well (Fig. 1I; Supplemental Fig. 1F). The lower number of B cells found in immunized mice - while a less profound effect than the reduction in GC - tempers the result, and it was possible that the function of helper CD4 T cells was impaired even their though numbers were not affected. Consistent with this hypothesis, the frequencies of Tfh and GC-Tfh cells were lower in *Dhrs7b*^Δ/Δ^ mice compared with controls (Supplemental Fig 1G-I). While not addressing whether or not B cells are part of the requirement for PexRAP in promoting GC, these results indicate that *Dhrs7b* is necessary for a normal-sized geminal center response.

**Figure 1.**
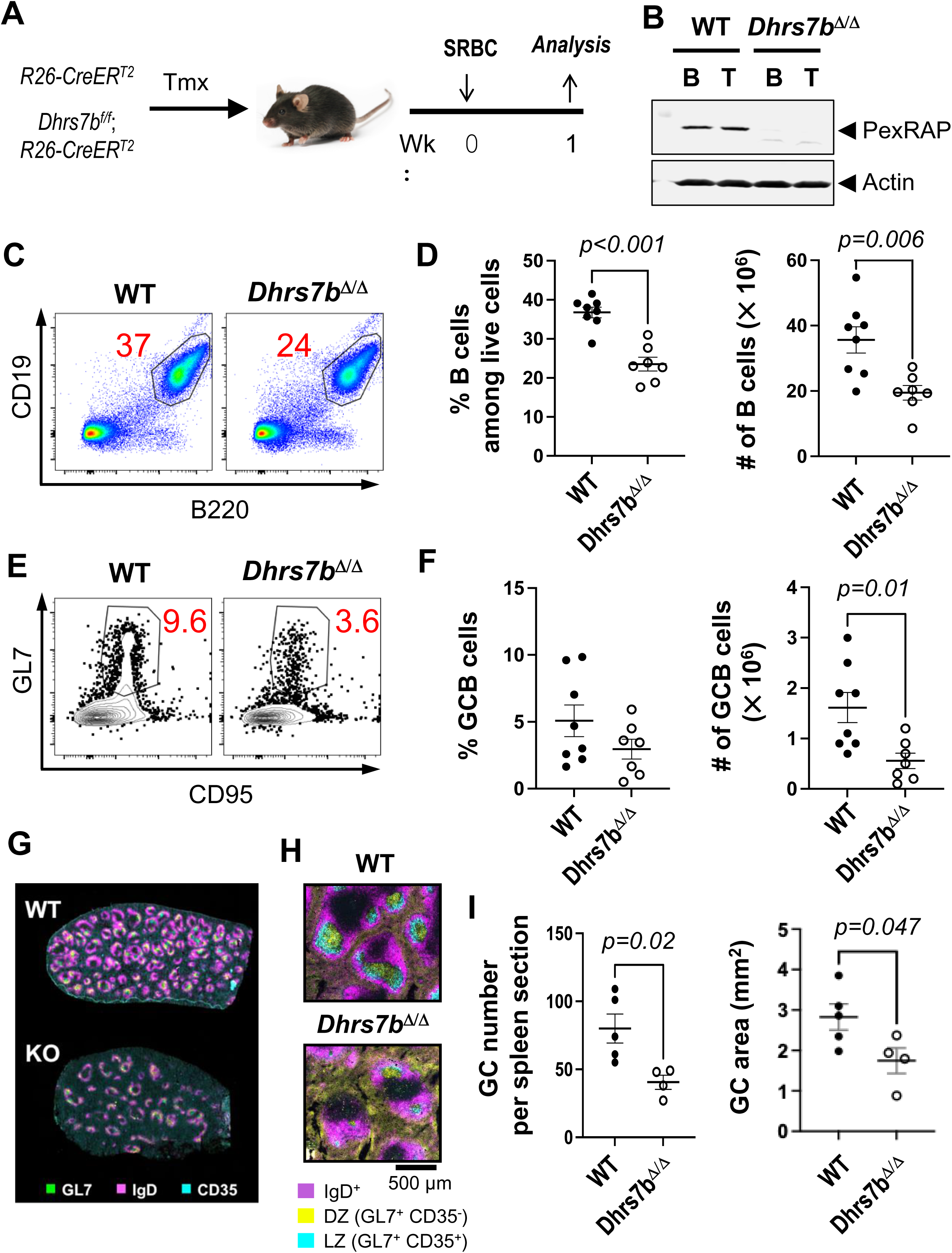
PexRAP promotes GC response. (A) Schematic of immunization with SRBC after inactivation of *Dhrs7b* in adult mice. Mice (*Rosa26*-CreER^T2^; *Dhrs7b^+/+^, or Rosa26*-CreER^T2^; *Dhrs7b^ff^*) were treated with tamoxifen, immunized with SRBC, and harvested 1 wk after immunization, as described in *Materials and Methods*. (B) Deletion efficiency of *Dhrs7b* conditional alleles. B and T lymphocytes were isolated from spleens of *Rosa26-*CreER^T2^ or *Dhrs7b^f/f^;Rosa26-*CreER^T2^ mice after in vivo tamoxifen injections followed by immunization with SRBC. WT and *Dhrs7b^Δ/Δ^*were analyzed by immunoblotting with antibodies directed against PexRAP (protein product of *Dhrs7b*) and actin (internal loading control). (C, D) PexRAP acutely regulates B cell numbers. Representative flow plots of viable splenic B cells (C) and aggregate data for three biologically independent replicate experiments (D) (n = 8 WT and 7 cKO). For data on T cells, see supplemental data Fig. S1A-C. (E, F) Effect of PexRAP on GC B cell response. Flow plots of GL7^+^ CD95^+^ GC B cells in the gate for viable IgD-negative (IgD^neg^), dump-negative B cells (E), and aggregated frequencies and numbers of GC B cells, as indicated, in the three replicate experiments (F). (G, H) PexRAP impact on GC response. After tamoxifen injections, *Dhrs7b* deficient mice [*Rosa26*-CreER^T2^; *Dhrs7b/^ff^* (46)] and WT controls (*Rosa26*-CreER^T2^) were immunized with SRBC and analyzed 7 d later, as in Fig. 1A-F. Data on gating strategy, GC counts per unit area and LZ area are in Supplemental Fig. 1D-F, and on Tfh cells in Supplemental Fig. 1G-I. (G, H) Shown are representative images from immunofluorescence staining of spleens with the indicated Ab in two independent experiments (5 WT vs 4 *Dhrs7b* cKO, i.e., *Dhrs7b*^Δ/Δ^), showing a low-power overview with many follicles (G) and a representative higher-magnification image to better delineate primary and secondary follicles (H). (I) Quantitation of number (left panel) and size (right panel) of GC in spleen sections, with GL7^+^, IgD^neg^ areas that include both CD35^+^ (LZ) and CD35^neg^ (DZ) areas. Mann-Whitney U test was used to calculate p values.

To determine if there are B lineage-specific functions of PexRAP, we used a conditionally active Cre transgene that is expressed specifically in mature B cells (67). *Dhrs7b* f/f, *huCD20-CreER^T2^*mice and *huCD20-CreER^T2^* were treated with tamoxifen to induce Cre-mediated recombination of the floxed alleles after initial establishment of normal B cell populations (Fig. 2A). Western blot analyses showed an almost complete absence of PexRAP protein from B cells purified from tamoxifen-treated *Dhrs7b^f/f^*; *huCD20-CreER^T2^* (hereinafter, *Dhrs7b*^Δ/Δ*-*B^) mice compared to similarly treated *huCD20-*CreER^T2^ controls (Fig. 2B). The enzyme encoded by this gene catalyzes the synthesis of a lipid precursor to many - but not all - plasmalogens and other ether phospholipids (46–48), and *Dhrs7b* deficiency impacted neutrophil survival (46). When we tested if *Dhrs7b* inactivation affected the steady-state B cell population, the frequencies and total numbers of CD19^+^ B220^+^ B cells in viable lymphocyte gates were about 20% lower in tamoxifen-treated *Dhrs7b*^Δ/Δ*-*B^ mice compared to tamoxifen-injected, *huCD20*-CreER^T2^, *Dhrs7b*^+/+^ controls (Fig. 2C). With loss-of-function after production of mature B cells, i.e., inactivation of *Dhrs7b* at ages over 6 wk, we observed balanced decreases in the numbers of multiple subsets within the B lineage without evidence of selectivity among main sub-classes of B cell. Moreover, the surface expression of IgM was comparable in *Dhrs7b*^Δ/Δ^ mice versus WT controls (Supplemental Fig 2. A-F). Thus, the frequencies of MZB and FOB amidst the smaller population of splenic B cells in tamoxifen-treated *Dhrs7b^f/f^*; *huCD20-CreER^T2^* mice were similar (Supplemental Fig 2B), and frequencies of B1 B cells in the peritoneal cavity also were unaffected (Supplemental Fig 2E). Prior work documents normal frequencies and numbers of developing and mature B cells under harsher conditions of tamoxifen-induced deletion with CreER^T2^ (65). De novo B cell production rates are too low to yield the ∼20% decrease observed one week after gene disruption, and few splenic B cells are in cell cycle. Accordingly, we infer that after maturation of a B cell, *Dhrs7b* promotes its survival, albeit to a modest degree.

**Figure 2.**
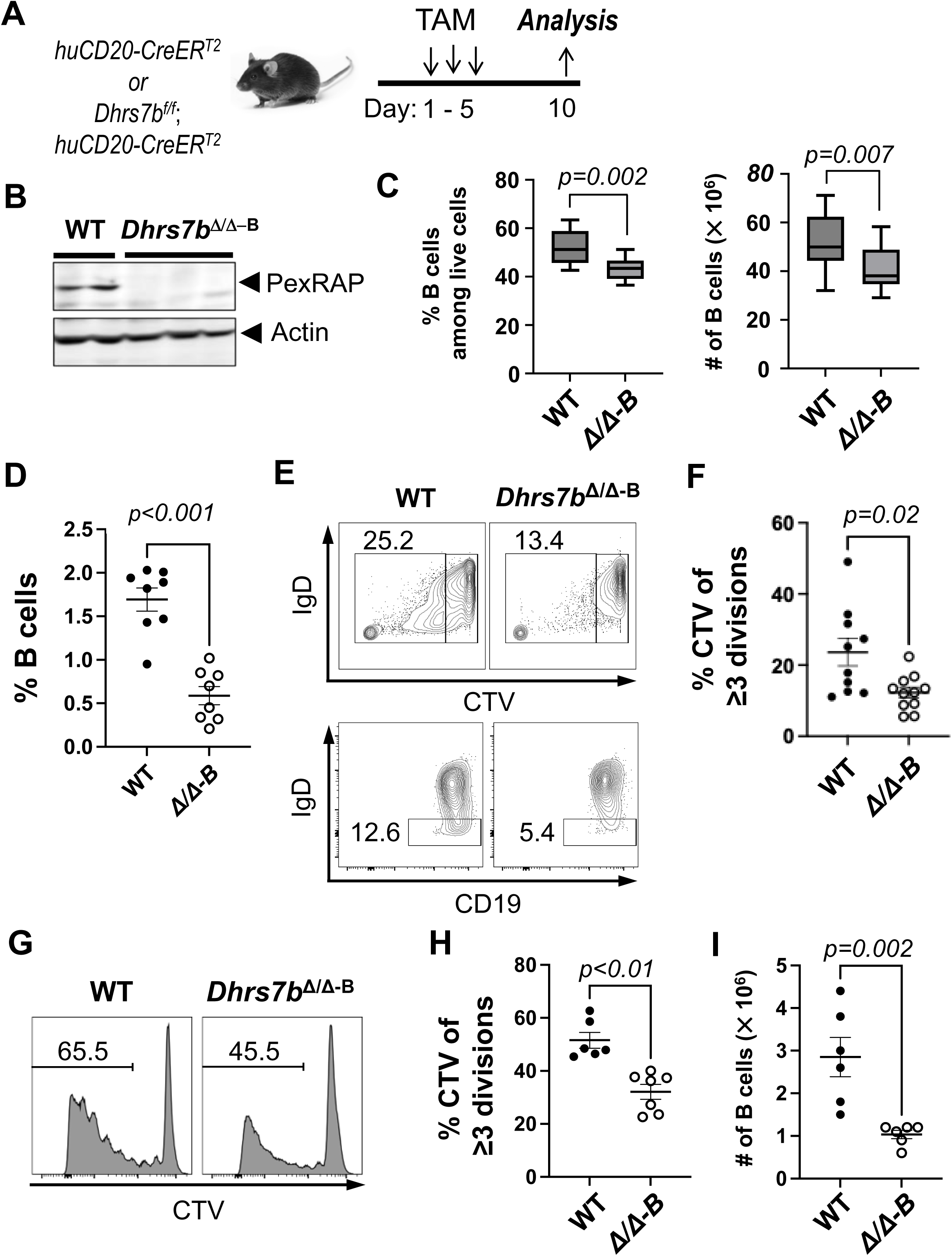
PexRAP promotes proliferation of B cells. (A) Schematic of *Dhrs7b* inactivation in mature mice via tamoxifen treatment and a B cell type-specific conditional allele. Mice (*huCD20*-CreER^T2^; *Dhrs7b^+/+^, or huCD20*-CreER^T2^; *Dhrs7b^f/f^*) were injected with tamoxifen (d1, 3, 5) and harvested at day 10. (B) Deletion efficiency of conditional *Dhrs7b* alleles. After in vivo tamoxifen injections, B cells were isolated from spleens of *huCD20-*CreER^T2^ or *Dhrs7b^f/f^; huCD20-*CreER^T2^ mice, as indicated, and analyzed by immunoblotting as in Fig. 1B. (C) PexRAP and the maintenance of B cells. Shown are the frequencies and the numbers of CD19^+^ B220^+^ B cells among viable lymphocytes in spleen (left and right panels, respectively). Data are pooled from four independent replicate experiments (n = 12 WT and 12 cKO). Shown in box and whisker plots are the means, with whiskers that extend to the minimum and maximum values and boxes that outline upper and lower quartile values with the midline identifying the median. (D-F) PexRAP regulates B cell proliferation in vivo. CTV-labeled B cells were adoptively transferred into µMT recipient mice and analyzed 4 days thereafter. Shown are the frequencies of B220^+^ CD19^+^ events among splenocytes in the viable cell gate (D), along with representative flow plots of CTV partitioning and surface IgD in B cell gates, with rectangles defining the gating for divided < or ≥ 3x, and IgD^+^ vs IgD^neg^. A similar difference was observed in analyzing divided vs undivided. (E). (F) Aggregated frequencies of divided B cells from five independent replicate experiments, as defined in (E) (n =10 WT and 11 cKO). Additional data on the IgD^neg^ population are in Supplemental Fig.2G. (G-I) PexRAP promotes B cell proliferation in vitro. After in vivo tamoxifen injections, bead-purified B cells from spleens of CreER^T2^ mice (WT and *Dhrs7b^Δ/Δ^, i.e., cKO*) were stained with Cell Trace Violet (CTV), activated and cultured 4 days in anti-CD40, BAFF, IL-4, IL-5 and 4-hydroxytamoxifen in three biologically independent experiments totaling 6 WT and 7 cKO mice). (G) Representative flow-cytometric analysis of CTV partitioning, with the gating line denoting multiply divided cells, with inset numbers representing the frequencies of such B cells in each of the two plots shown. (H) Quantified frequencies of divided ≥3 times. (I) Numbers of B cells recovered at the end of the cultures.

B cell population growth can be impacted by increased cell death and/or decreased proliferation. To test if PexRAP affects proliferation in vivo, *Dhrs7b*^Δ/Δ^ B cells were stained with CellTrace Violet (CTV) and adoptively transferred into B cell deficient µMT recipient mice. Strikingly fewer PexRAP-depleted B cells were recovered (Fig. 2D), and the frequencies of *Dhrs7b*^Δ/Δ^ B cells that were IgD^neg^ (i.e., had been activated) were half those of controls (Fig. 2E; Supplemental Fig. 2G). As compared to control B cells, CTV partitioning analyses suggested that division rates in vivo were modestly lower for the *Dhrs7b*^Δ/Δ^ B cells (Fig. 2E, F). Mitogen-stimulated *Dhrs7b*^Δ/Δ^ B cells that proliferated in culture exhibited substantially less robust division: frequencies of viable cells that had undergone ≥3 divisions were halved in *Dhrs7b*^Δ/Δ^ B cells) and yielded ∼1/3 as large a progeny population (Fig. 2G-I; Supplemental Fig 2H). Collectively, these data indicate that the *Dhrs7b* gene product supports homeostatic maintenance of a quiescent (pre-immune) B cell population *in vivo* and effective proliferation of B cells.

### B cell expression of PexRAP is required for achieving normal concentrations of ether phospholipids

As noted, lipidomic analyses that used 2-dimensional image mass-spectrometry (2D-IMS) discovered that local concentrations of a subset of ether lipids are enriched in GC compared with the area outside of GC after immunization (45). To measure the extent to which PexRAP expression within mature B cells can affect their lipid content and composition, including their ether- and plasmalogen phospholipids, LC-MS-MS analyses were performed with B cells directly isolated from spleens as well as those cultured after mitogenic activation. These analyses showed substantial reductions in many ether phospholipids - as well as lysophosphatidylethanolamines (LPE) generated by phospholipase cleavage of the R2 fatty acid sidechain of an ether phospholipid - in PexRAP-deficient B cells (Fig. 3A, B; Supplemental Fig. 3A, B). While beyond the scope of this work, this evidence suggests that generation of potential signaling molecules via phospholipase(s) such as PLA2 may be reduced. Of note, *Dhrs7b* inactivation reduced levels of a subset of plasmalogens in naive B cells (Fig. 3A, B, D; Supplemental Fig 3A) though changes of others were modest or absent in the naive population. A greater impact of PexRAP on quantitative amounts of phospholipids was found in the activated B cells. Several species, skewed towards those with the most detected ions, a number of which were at less than 1/5th the level of non-deleted control B cells (Fig. 3C, D). Phospholipid results observed with naive and activated B cells from tamoxifen-treated CreER^T2^-expressing mice were similar to those of unmanipulated controls (Supplemental Fig. 3C, D). There is no in vitro surrogate that faithfully represents GC or the GC B cell, but collectively these data provide strong evidence that the capacity to rapidly produce increased levels of many ether lipid molecules after lymphocyte activation depends on a functional *Dhrs7b* gene in B cells. The data provide evidence that *Dhrs7b* in B cells enhances cell-autonomous biosynthesis of some plasmalogens and other ether lipid species. Especially in resting B cells that are out of cycle, additional uptake or biosynthesis pathways may add to the overall pool (46, 47). Notwithstanding these potential contributions, however, the straightforward inference is that PexRAP catalyzes the generation of ether lipid precursors in amounts crucial for regulating the overall ether phospholipid pools, especially after B cell activation.

**Figure 3.**
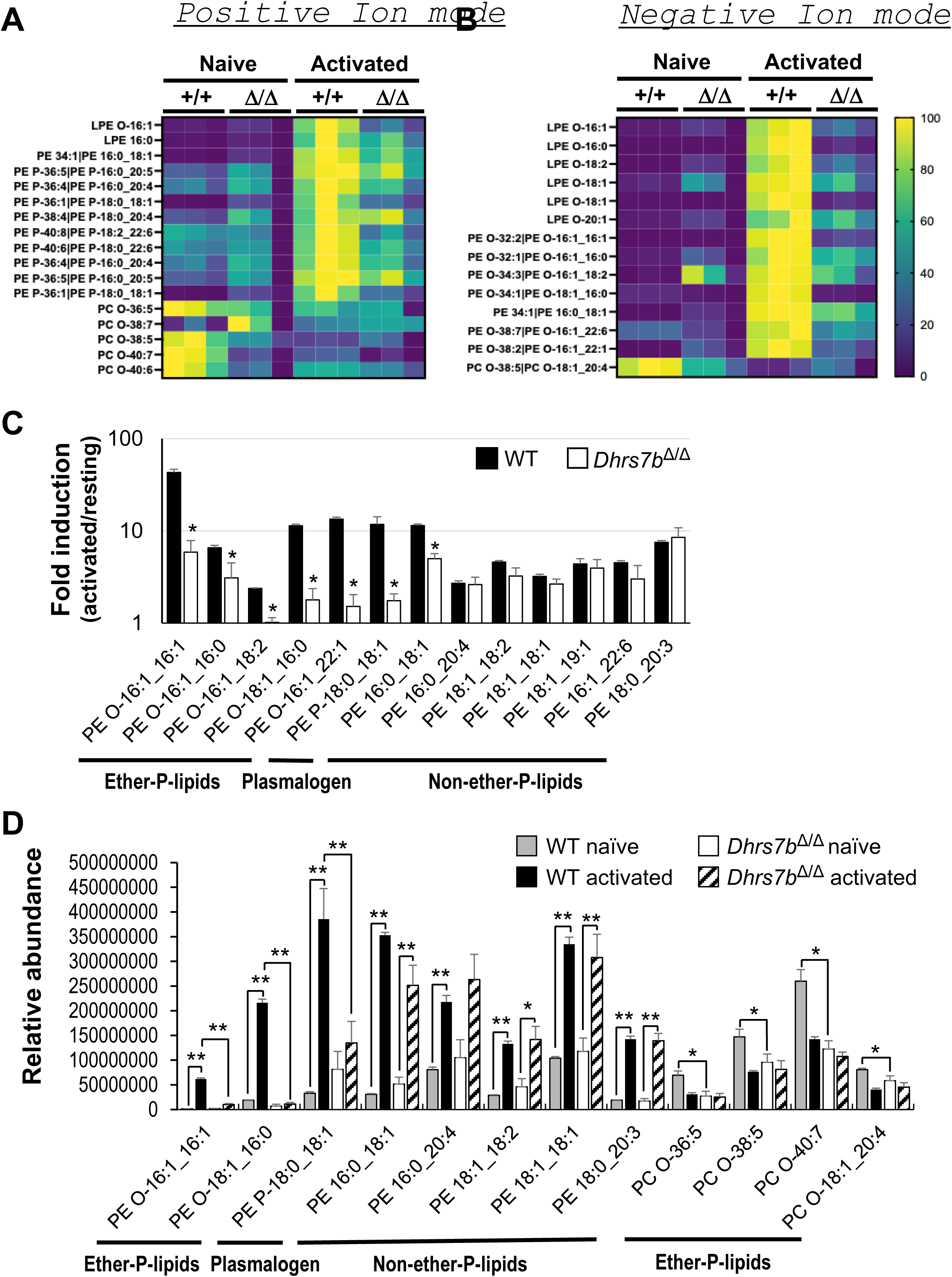
B cell expression of PexRAP is essential for normal concentrations of many ether phospholipids. Splenic B cells (CreER^T2+^ or CreER^T2+^, *Dhrs7b* Δ/Δ) from tamoxifen-injected mice were analyzed by LC-MS-MS either directly ex vivo (“Naive”) or after activation and culture (48 h) (“Activated”) as in Fig. 2. Shown are z-scored heat maps for the relative abundance of subsets of ether and plasmalogen phospholipids and lysophospholipids identified by exact mass and secondary fragmentation in (A) positive and/or (B) negative ion modes. PE, phosphatidylethanolamine; PC, phosphatidylcholine; LPE, lysophosphatidylethanolamine. More species are displayed in Supplemental Fig. 3. (C) Quantitated peak areas for activation-induced increases (’fold-induction’ ratios of activated/resting, plotted on log_10_ scale) for the indicated ether, plasmalogen, and diacyl (non-ether) phospholipids in the PexRAP-sufficient and -depleted B cells. * denotes species for which P<0.05 for the effect of *Dhrs7b* inactivation. (D) Shown are the raw values of mean peak areas for the indicated species in the freshly purified and the activated B cells of the indicated genotypes, as indicated. (Three independent samples from individual mice of each genotype were analyzed.) * p<0.05, ** p<0.01 by unpaired Student’s. t-test. Additional data including controls comparing B cells of unmanipulated B6 mice to those expressing CreER^T2^ and injected with tamoxifen are in Supplemental Fig. 3.

Our earlier work (45) reported only ion mode features in negative mode and did not analyze the primary follicle (the vast majority of which are resting B cells) relative to the extrafollicular white pulp, red pulp, and GC. To investigate the requirement for specific expression of PexRAP, we started with AID-GFP mice to enhance spatial localization. These measurements confirmed that a substantial number of both positive and negatively charged ions whose exact masses identified them as plasmalogens were substantially concentrated in GC as compared to the rest of the B cell follicle (Table 1; Supplemental Fig. 4A, B). Accordingly, we tested if the differential enrichment of any ether lipid species in primary or secondary follicles depends on biosynthesis within a specific lymphoid lineage. Mice with conditional mutations were tamoxifen-treated, immunized with SRBC, harvested seven days thereafter, and analyzed by IMS (Fig. 4A). With deletion driven by the widely expressed *Rosa26-CreER^T2^* transgene, as in earlier work (46) and Fig. 1, substantial reductions of the GC enrichment pattern were observed (Fig. 4B). For instance, two representative ions identified in negative ion mode (i.e. *m/z* 752.5545 and *m/z* 776.5556) were again enriched in GC regions of spleens from immunized WT mice. These features barely increased in splenic samples of immunized *Dhrs7b^Δ/Δ^* (tamoxifen-injected *Dhrs7b*^f/f^; *Rosa26-CreER^T2^)* mice (Fig. 4B).

**Table 1.** Immunization-induced increases of selected lipids in GC.

| **Negative Ions** |  |  |  |
| --- | --- | --- | --- |
| **m/z<sup>a</sup>**<br>**Lipid ID** | **UI** | **1° B cell Follicle** | **GC** |
| **716.5216**<br>PE(18:0_16:1) | 618 ± 22<sup>b</sup> | 2400 ± 83<sup>c</sup> | 3674 ± 154<sup>d</sup> |
| **752.5545**<br>PE(O-18:0_20:4) | 301 ± 14 | 1739 ± 33<sup>c</sup> | 1960 ± 52<sup>d</sup> |
| **760.5056**<br>PE(16:0_22:6) | 203 ± 4 | 1816 ± 30<sup>c</sup> | 2031 ± 23<sup>d</sup> |
| **772.5242**<br>PE(P-18:1_22:6) | 234 ± 5 | 2334 ± 47<sup>c</sup> | 2578 ± 63<sup>d</sup> |
| **776.5556**<br>PE(O-18:0_22:6) | 277 ± 9 | 1638 ± 51<sup>c</sup> | 2074 ± 90<sup>d</sup> |
| **869.5526**<br>PI(O-18:0_20:5) | 242 ± 7 | 1611 ± 24<sup>c</sup> | 1789 ± 15<sup>d</sup> |
| **872.576**<br>PC(18:0_24:0) | 171 ± 6 | 1167 ± 16<sup>c</sup> | 1369 ± 18<sup>d</sup> |
| **888.5612**<br>PS(22:1_22:6) | 506 ± 13 | 2684 ± 54<sup>c</sup> | 3386 ± 72<sup>d</sup> |

| **Positive Ions** |  |  |  |
| --- | --- | --- | --- |
| **m/z<sup>a</sup>**<br>**Lipid ID** | **UI** | **1° B cell Follicle** | **GC** |
| **739.4587**<br>PG(12:0_20:3) | 3042 ± 102 | 3733 ± 116<sup>c</sup> | 4778 ± 274<sup>d</sup> |
| **740.465**<br>PC(16:0_15:1) | 1310 ± 13 | 1478 ± 66<sup>c</sup> | 1933 ± 142<sup>d</sup> |
| **784.5564**<br>PC(10:0_24:0) | 2840 ± 24 | 7956 ± 147<sup>c</sup> | 9856 ± 270<sup>d</sup> |
| **798.5347**<br>PC(10:0_25:0) | 9937 ± 269 | 27589 ± 574<sup>c</sup> | 34733 ± 1644<sup>d</sup> |
| **799.5375**<br>PG(18:0_18:1) | 5665 ± 65 | 12078 ± 282<sup>c</sup> | 15311 ± 790<sup>d</sup> |

**Figure 4.**
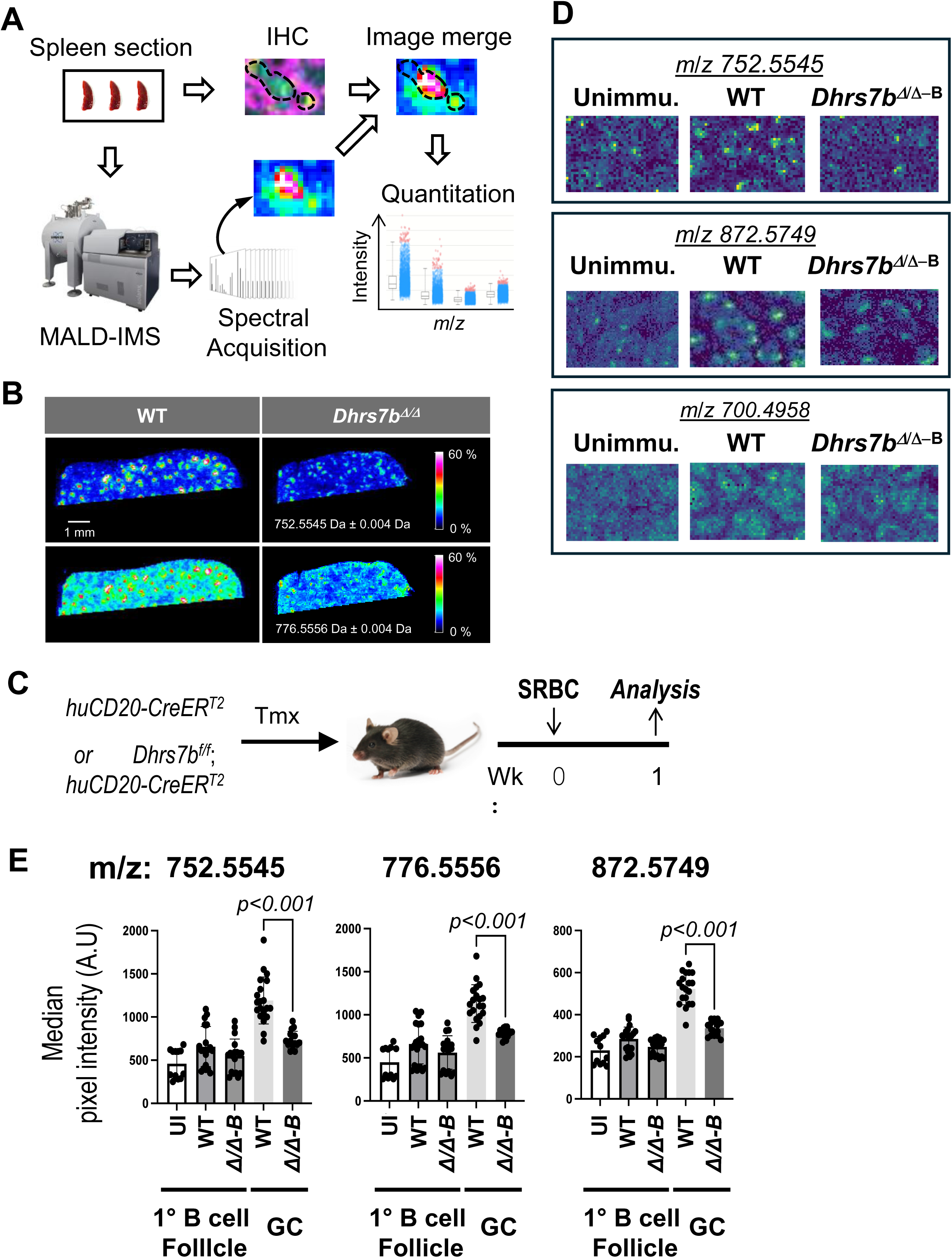
PexRAP is essential for normal concentrations and distributions of some ether phospholipids in splenic follicles. (A) Schematic diagram illustrating the 2D-IMS analysis work flow, image merging and quantitation. Spleens harvested 1 wk after immunization with SRBC were used to generate serial tissue sections followed by immunofluorescence (IF) staining of one section and IMS analysis with the adjacent one. IF and IMS images were aligned to map ion intensity distributions to microanatomic regions (B cell follicle and GC). The intensities of specific ions on B cell follicles and GC regions were quantitated as described in *Materials and Methods.* (B) Identification of ether lipid species localizing to lymphoid follicles. Representative ion images of two ions [*m/z* 752.5545, and *m/z* 776.5556] with spleens from immunized mice (WT and *Dhrs7b^Δ/Δ^*) as shown in Fig. 1A. (C) Schematic of immunization for IMS analyses of B cell-specific PexRAP loss. Mice of the indicated genotypes (huCD20-CreER^T2^ ± *Dhrs7b* f/f) were treated with tamoxifen, immunized with SRBC, and harvested 1 wk after immunization. (D) Identification of ether lipid species localizing to lymphoid follicles. Representative ion images of three ions [*m/z* 752.5545, *m/z* 872.5749, and *m/z* 700.4958] in mass spectrometry imaging of spleens from WT and *Dhrs7b^Δ/Δ−B^* mice. Immunofluorescent images at higher magnification, delineating LZ and DZ marked by CD35 staining, along with quantification of sizes of GC and their LZ and DZ, are shown in Supplemental Fig. 5A, B. (E) Shown in the bar graphs are the mean (±SEM) ion intensities in primary lymphoid follicles (B cell zones) and GC from spleens of WT and *Dhrs7b^Δ/Δ-B^*mice, immunized or not (“UI”) as indicated. Median intensity of each ion was obtained from 3 follicles / spleen and 3 GC / spleen from WT and *Dhrs7b^Δ/Δ-B^* mice (three biological replications comprising 7 WT and 6 cKO spleens). P values were calculated by Mann-Whitney U test.

To determine if the levels of particular ether phospholipids in GC regions were affected by PexRAP expression in B cells, *Dhrs7b^f/f^*; *huCD20-*CreER^T2^ mice and *huCD20-*CreER^T2^ controls were analyzed after tamoxifen injections followed by immunization, and compared to samples from unimmunized mice (Fig 4C). IMS using both positive and negative ion modes identified at least eight ions - including *m/z* 752.5545, *m/z* 776.5556, and *m/z* 872.5749 - much more substantially (∼2-3-fold) concentrated in GC of immunized control mice (Fig. 4D, E; Table 2) than in the primary follicle of unimmunized mice. Notably, these increased signals in secondary follicles (GC) were reduced or almost completely eliminated in GCs of mice immunized after B cell-specific PexRAP depletion (*Dhrs7b*^Δ/Δ-B^) (Fig. 4D, E; Table 2; Supplemental Fig. 4C). Collectively, these results indicate that *Dhrs7b* gene expression in activated B cells regulates the spectrum of ether lipids in the micro-anatomic locale of a secondary follicle. Interestingly, immunization also led to modest increases in the levels of some ether and conventional phospholipids in the B cell-rich primary follicle regions (Table 2). This finding extends the previous report (45), which focused on the relationship between the AID-GFP^hi^ region (i.e., GC) and lipid features. Moreover, the absence of PexRAP blunted most of the immunization-induced increases observed in the primary follicles. These findings and those of Fig. 3 suggest that there are secondary consequences of *Dhrs7b* inactivation (e.g., some di-acyl phospholipids are affected), but provide direct evidence that the enhancement of many ether phospholipid signals in the GC depends on PexRAP in B cells.

**Table 2.** Impact of PexRAP on relative quantities of selected lipid species in primary and secondary follicles (GC).

| Negative Ions |  |  |  |  |  |
| --- | --- | --- | --- | --- | --- |
| m/z<sup>a</sup><br>Lipid ID | UI | WT follicle | PexRap cKO follicle | WT GC | PexRap cKO GC |
| **436.2801**<br>LPE (O-16:1) | 315 ± 4 | 369 ± 7 | 374 ± 11 | 610 ± 20<sup>e</sup> | 475 ± 19 |
| **716.5216**<br>PE (18:0_16:1) | 754 ± 43<sup>b</sup> | 1608 ± 182<sup>c</sup> | 1263 ± 112 | 2729 ± 247<sup>e</sup> | 1770 ± 137<sup>f</sup> |
| **746.5098**<br>PE (O-16:1_22:6) | 622 ± 19 | 672 ± 35 | 609 ± 23 | 1066 ± 39<sup>e</sup> | 836 ± 39 |
| **752.5545**<br>PE (O-18:0_20:4) | 459 ± 49 | 657 ± 51<sup>c</sup> | 548 ± 46<sup>d</sup> | 1191 ± 61<sup>e</sup> | 734 ± 25<sup>f</sup> |
| **760.5056**<br>PE (16:0_22:6) | 229 ± 8 | 386 ± 25<sup>c</sup> | 312 ± 16<sup>d</sup> | 569 ± 27<sup>e</sup> | 408 ± 15<sup>f</sup> |
| **772.5242**<br>PE (P-18:1_22:6) | 388 ± 47 | 617 ± 46<sup>c</sup> | 499 ± 33<sup>d</sup> | 918 ± 21<sup>e</sup> | 646 ± 9<sup>f</sup> |
| **776.5556**<br>PE (O-18:0_22:6) | 449 ± 53 | 661 ± 50<sup>c</sup> | 562 ± 46 | 1130 ± 49<sup>e</sup> | 789 ± 13<sup>f</sup> |
| **869.5526**<br>PI (O-18:0_20:5) | 509 ± 81 | 641 ± 50<sup>c</sup> | 494 ± 40<sup>d</sup> | 1014 ± 21<sup>e</sup> | 690 ± 12<sup>f</sup> |
| **872.576**<br>PC (18:0_24:0) | 230 ± 19 | 285 ± 12<sup>c</sup> | 247 ± 9<sup>d</sup> | 522 ± 17<sup>e</sup> | 336 ± 9<sup>f</sup> |
| **888.5612**<br>PS (22:1_22:6) | 1423 ± 277 | 2051 ± 199<sup>c</sup> | 1904 ± 199 | 3260 ± 156<sup>e</sup> | 2629 ± 80<sup>f</sup> |

| Positive Ions |  |  |  |  |  |
| --- | --- | --- | --- | --- | --- |
| m/z<sup>a</sup><br>Lipid ID | UI | WT follicle | PexRap cKO follicle | WT GC | PexRap cKO GC |
| **730.5767**<br>PE (P-18:0_18:1) | 91 ± 3 | 97 ± 3 | 85 ± 3 | 130 ± 7<sup>e</sup> | 102 ± 4 |
| **739.4587**<br>PG (12:0_20:3) | 3042 ± 102 | 3094 ± 40 | 2944 ± 31<sup>d</sup> | 3506 ± 140<sup>e</sup> | 3011 ± 46<sup>f</sup> |
| **784.5564**<br>PC (10:0_24:0) | 2840 ± 24 | 2917 ± 74 | 2617 ± 46<sup>d</sup> | 3522 ± 102<sup>e</sup> | 2561 ± 42<sup>f</sup> |
| **798.5347**<br>PC (10:0_25:0) | 9937 ± 269 | 10928 ± 214<sup>c</sup> | 11372 ± 155<sup>d</sup> | 15378 ± 116<sup>e</sup> | 14694 ± 186<sup>f</sup> |
| **801.5509**<br>PG (18:0_18:0) | 1540 ± 37 | 1591 ± 46 | 1633 ± 26 | 1444 ± 27<sup>e</sup> | 1533 ± 22<sup>f</sup> |
| **847.5361**<br>PI (15:0_18:0) | 1277 ± 24 | 1339 ± 54 | 1217 ± 35 | 1150 ± 46<sup>e</sup> | 1044 ± 44 |
| **874.5623**<br>PC (20:5_22:6) | 754 ± 39 | 688 ± 48 | 675 ± 25 | 469 ± 27<sup>e</sup> | 556 ± 37 |

### B cell-intrinsic role of *Dhrs7b* in Ab affinity and quantity

As key components of adaptive immunity, progeny of an activated B cell can differentiate into Ab-secreting plasma cells (PC) or into GC B cells. Over the course of an immune response, the GC reaction diversifies the affinity for and breadth of antigen recognized by the original BCR, and improves properties of memory [(14, 17–19, 68). PC that develop after a second encounter with antigen increase the amounts of high-affinity Ab. The higher quality and quantity of Ag-specific Abs from plasma cells are integral to humoral immune responses (68). An increasing body of evidence indicates that metabolic reprogramming and intermediary metabolites modulate immune cell differentiation and function (26–36). The observed B cell-intrinsic function of *Dhrs7b* in the accumulation of many ether lipid species in both primary and secondary follicles prompted us to test the effect of B cell type-restricted PexRAP depletion on GC responses and Ag-specific Ab production. Tamoxifen-injected *Dhrs7b^f/f^*; *huCD20-CreER^T2^*and control mice were immunized with NP-ovalbumin (NP-OVA), then boosted with NP-OVA to elicit affinity-matured Ab, and harvested 1 week thereafter (Fig. 5A). Although PexRAP-deficient B cell numbers were reduced less than 20% a week after starting gene inactivation prior to immunization (i.e., were over 0.8-fold those of controls) (Fig 2A-C), frequencies and numbers of IgD^neg^ B cells were halved in *Dhrs7b*^Δ/Δ-B^ mice at harvest (4 wk after completion of the induced deletion) (Fig. 5B, C). Moreover, the GL7^+^ CD95^+^ fraction of the B cell population in the dump / IgD^neg^ gate was substantially reduced (Fig. 5D), such that numbers of GL7^+^ CD95^+^ GC B cells were dramatically decreased in *Dhrs7b*^Δ/Δ-B^ mice (Fig. 5E). Consistent with these findings, GC in *Dhrs7b*^Δ/Δ-B^ mice were reduced after SRBC immunization (Fig 5 F, G; Supplemental Fig. 5A, B), and the numbers of centrocytes and centroblasts were dramatically lower in *Dhrs7b*^Δ/Δ-B^ mice compared with WT controls (Fig 5H). LZ B cells (CD86^+^ CXCR4^lo^) trended toward being at lower prevalence in *Dhrs7b*^Δ/Δ-B^ mice, whereas their DZ counterparts (CD86^neg^ CXCR4^+^) were comparable to wildtype controls, resulting in a modest decrease in the LZ/DZ ratio when B cells lacked PexRAP (Supplemental Fig. 5C). The frequencies of CD138^+^ GL7^+^ IgD^-^early plasmablasts were comparable between *Dhrs7b*^Δ/Δ-B^ mice and control mice (Supplemental Fig 5D). Although the loss-of-function was B cell specific, the frequencies of Tfh and GC-Tfh cells were decreased in *Dhrs7b*^Δ/Δ-B^ mice compared with control mice (Supplemental Fig 5E). We infer that support for the Tfh populations was reduced due to impact(s) on GC B cell numbers and / or function.

**Figure 5.**
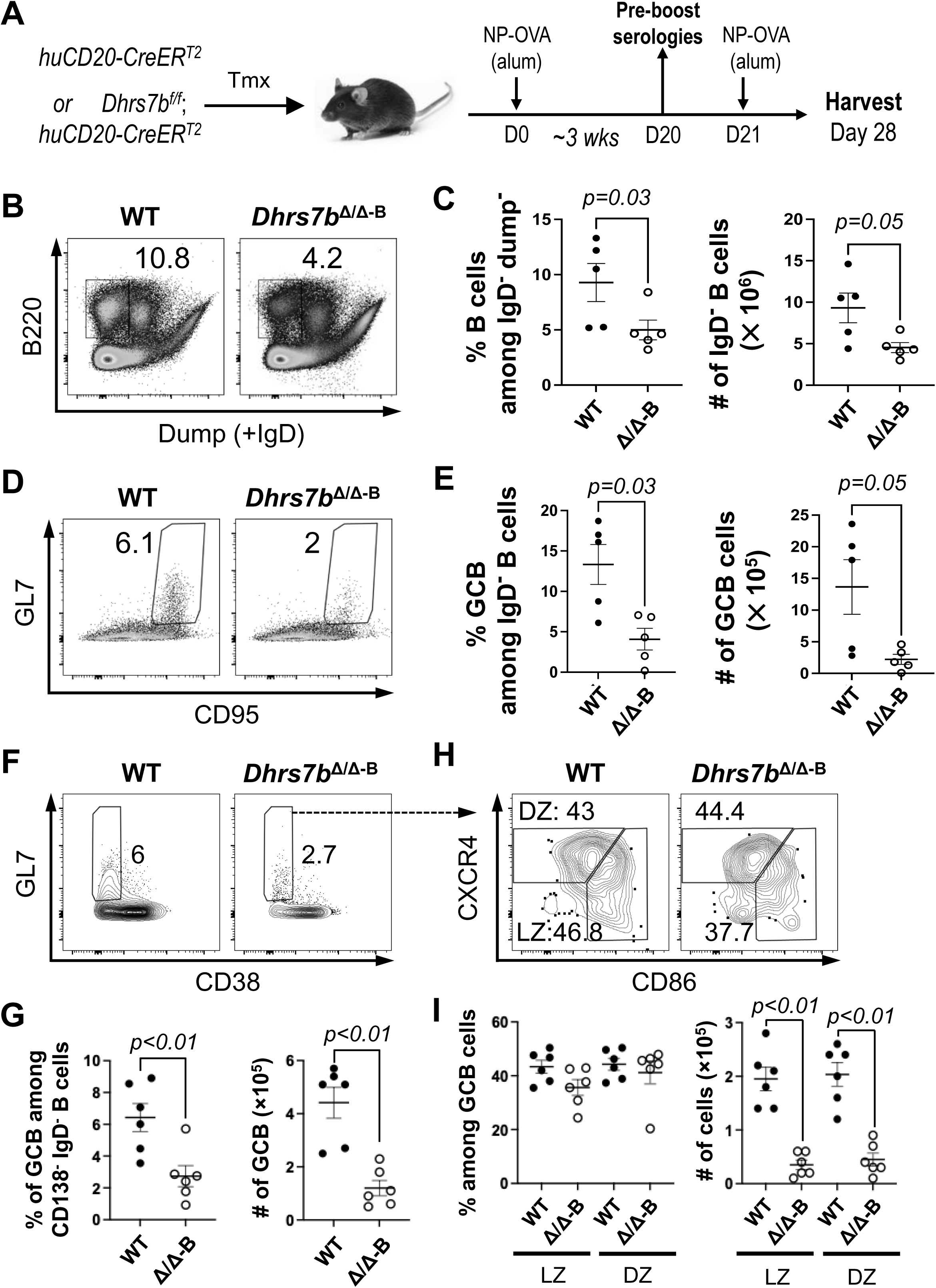
B cell intrinsic role of PexRAP in GC response. (A) Schematic of immunization with NP-OVA in alum after inactivation of *Dhrs7b* in B lineage cells. Tamoxifen-treated WT (*huCD20*-CreER^T2^; *Dhrs7b^+/+^*) or *Dhrs7b^Δ/Δ-B^* mice *(huCD20*-CreER^T2^; *Dhrs7b^ff^*) were immunized with NP-OVA, with sera collected 3 wk thereafter (“1^0^ response”), followed by boosting with NP-OVA and harvest 1 wk after the 2nd immunization. (B, C) Representative flow plots of splenic IgD^neg^ B cells (B) at the time of harvest (1 wk after 2nd immunization), and aggregate data (C) for two replicate experiments (n = 5 WT and 5 cKO). The cocktail of reagents for the dump channel included anti-IgD. (D, E) B cell-intrinsic function of PexRAP in GC B cell response. Representative flow plots of GL7^+^ CD95^+^ GC B cells among viable IgD^neg^ B cells (D), and aggregated frequencies and numbers of GC B cells for two replicate experiments (E). Mann-Whitney U test was used to calculate p values. (F, G) WT (*huCD20*-CreER^T2^; *Dhrs7b^+/+^*) or *Dhrs7b^Δ/Δ-B^* mice *(huCD20*-CreER^T2^; *Dhrs7b^ff^*) were injected with tamoxifen and immunized with SRBC as in Fig 4A. Shown are the representative flow plots of GL7^+^ CD38^-^GC B cells among IgD^neg^ CD138^neg^ viable B cells (F), and aggregate data (G) for two replicate experiments (n = 6 WT and 6 cKO). (H) PexRAP is mostly dispensable for the balance of LZ and DZ B cells. The graph shows the mean (± SEM) frequencies (left) and the numbers (right) of CD86^+^ CXCR4^lo^ LZ B cells and CD86^neg^ CXCR4^+^ DZ B cells among IgD^neg^ CD38^neg^ CD138^neg^ GL7^+^ GC cells. Mann-Whitney U test was used to calculate p values.

Ab class-switch recombination (CSR) and affinity maturation can occur through extrafollicular responses. However, the GC reaction significantly increases Ab diversification and high-affinity Ab production, in part later in a primary response but also upon secondary exposure to antigens (6, 8, 9, 17–19). ELISA performed with sera at 3 wk post-immunization (Fig. 6A-D), just before the boost, showed that B cell-specific depletion of PexRAP prior to immunization reduced NP-specific IgM and the switched isotype IgG1 (Fig. 6A, C). Of note, the capacity to generate high-affinity Ab, detected with low valency NP_2_, was even more severely undermined than the overall response - especially for IgG1 (Fig. 6B, D). A week after a second immunization - an interval similar to that which followed immunization with SRBC - Ab-secreting cells (ASCs) in the spleen (Fig. 6E) and circulating anti-NP IgM concentrations were diminished in *Dhrs7b*^Δ/Δ-B^ mice compared with controls (*huCD20*-CreER^T2^, *Dhrs7b*^+/+^ treated with tamoxifen in parallel to the mice with induced loss of PexRAP) (Fig. 6F). The ratio of high-affinity (NP_2_-binding) to all-affinity (NP_20_) IgM Ab also was substantially lower in *Dhrs7b*^Δ/Δ-B^ mice (Fig. 6G), as were the serum levels of high-affinity IgG1 and IgG2c, class-switched isotypes (Supplemental Fig. 5F, G). These data reinforce and extend the conclusion that the expression of PexRAP in mature B cells is important for their physiology and function.

**Figure 6.**
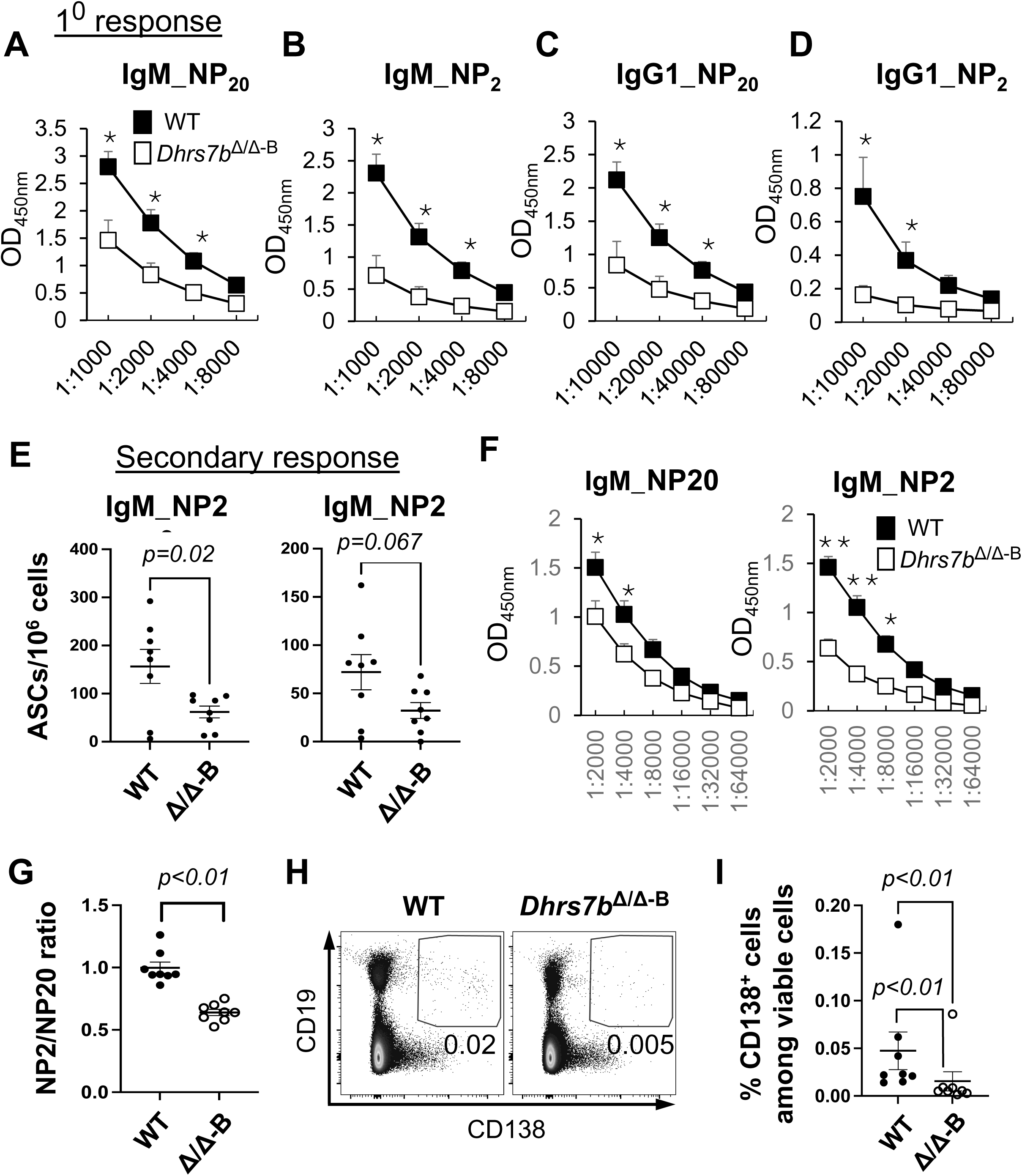
Ab response and affinity increase promoted by PexRAP in B cells. Tamoxifen-treated mice (*huCD20*-CreER^T2^; *Dhrs7b^+/+^* or *huCD20*-CreER^T2^; *Dhrs7b^f/f^*, i.e., *Dhrs7b^Δ/Δ-B^*) were immunized as in Fig. 5, with venous blood collected to measure Ag-specific Ab in the 1^0^ response just prior to a second immunization, followed by harvesting a week thereafter (2^0^ response). (A-D) Ag-specific Ab in primary response sera, prior to the boost. Shown are the all-(NP_20_) and high-(NP_2_)-affinity anti-NP IgM (A, B) and IgG1 (C, D), as indicated, with NP_20_ a high hapten density to detect both low- and high-affinity Ab and NP_2_ a low hapten density selective for high-affinity Ab. (E, F) Levels of anti-NP IgM detected using NP_20_ and NP_2_ for ELISpot (E) and ELISA (F) as described in *Materials and Methods*. Graphs show mean (± SEM) number of Ab-secreting cells in spleen (E) and (F) mean (± SEM) OD_450nm_ in measurement of NP-specific IgM in serial dilutions of sera. P values were calculated by students’ t-test. * indicates p<0.05, and ** indicates p<0.01. (G) PexRAP in B cells promotes Ab affinity maturation. The bar graph shows the mean (± SEM) ratios of high-affinity to all-affinity NP-specific IgM Ab in sera of individual mice (each dot representing one subject) using OD_450nm_ values at the 1:2000 dilution, with data from three independent experiments comprising 8 mice of each type (WT; *Dhrs7b^Δ/Δ−B^*). P values were calculated by Mann-Whitney U test. (H, I) PexRAP promotes CD138^+^ cell differentiation in vivo. B cells were adoptively transferred into µMT recipient mice and analyzed at 4 days after transfer. Representative flow plot of CD138 and cD19 expression (H) and aggregated frequencies of CD138^+^ CD19^+^ cells in the viable cell gate (I) from four independent replicate experiments. To test for a potential distortion arising from outlier values, statistical testing was performed both with and without their inclusion.

Consistent with *in vivo* results finding reduced Ag-specific ASCs in *Dhrs7b*^Δ/Δ-B^ mice compared with controls (Fig. 6E), the attenuated population of PexRAP-deficient B lineage cells recovered after transfer into recipient mice (Fig. 2D) yielded lower frequencies of CD138^+^ progeny (Fig. 6H, I) along with evidence of reduced activation [i.e., lower frequencies of IgD^-^ progeny (Supplemental Fig. 2G). Mitogen-stimulated *Dhrs7b*^Δ/Δ^ B cells also yielded lower frequencies of CD138^+^ cells in vitro, but the division-specific frequencies of CD138^+^ cells showed only modest decreases in *Dhrs7b*^Δ/Δ^ B cells (Supplemental Fig. 6 A, B). Thus, the reduction of CD138^+^ cell differentiation appears to be due mostly to its dependence on survival to a sufficient division count.

*Dhrs7b* inactivation was initiated prior to immunization in the preceding experiments. Therefore, the impact of PexRAP deficiency on GC and the Ab response in such a setting might be due exclusively to impairment of B cells, e.g., their reduced population expansion, prior to their entry into GC. Alternatively, the effects could also in part involve a requirement for PexRAP-dependent metabolites within GC B cells. To test if *Dhrs7b* functions within GC, we used conditional deletion of *Dhrs7b* driven by the *S1pr2-CreER^T2^* transgene whose expression at high levels marks GC B cells (69). Of note, experiments with a fate-marking reporter allele showed that activated B cells that lack the GC B phenotype were not marked by this conditional Cre after immunization with the NP-carrier approach (69). *Dhrs7b^f/f^*; *S1pr2-CreER^T2^* and control *S1pr2-CreER^T2^* (control) mice were immunized with SRBC, and tamoxifen was injected at a time point after the initiation of GC to test more directly that the inactivation of *Dhrs7b* would be in GC B cells rather than pre-GC blasts (Figure 7A). Frequencies of GC B cells were substantially lower in tamoxifen-treated *Dhrs7b^f/f^*; *S1pr2-*CreER^T2^ mice (Figure 7B, C). While this finding does not exclude that there may be an additional effect of impaired clonal expansion in B cells activated by immunization but not yet resident in the GC, the results indicate that PexRAP functions within GC B cells to support a full population of this subset. All together, these data provide evidence that *Dhrs7b* expression influences the lipidome and function of GC B cells. Notably, an optimal GC response requires PexRAP function in B cells, and the B cell-intrinsic functions of *Dhrs7b* promote the quantity and affinity maturation of Ab responses, in part via net proliferation of B cells.

**Figure 7.**
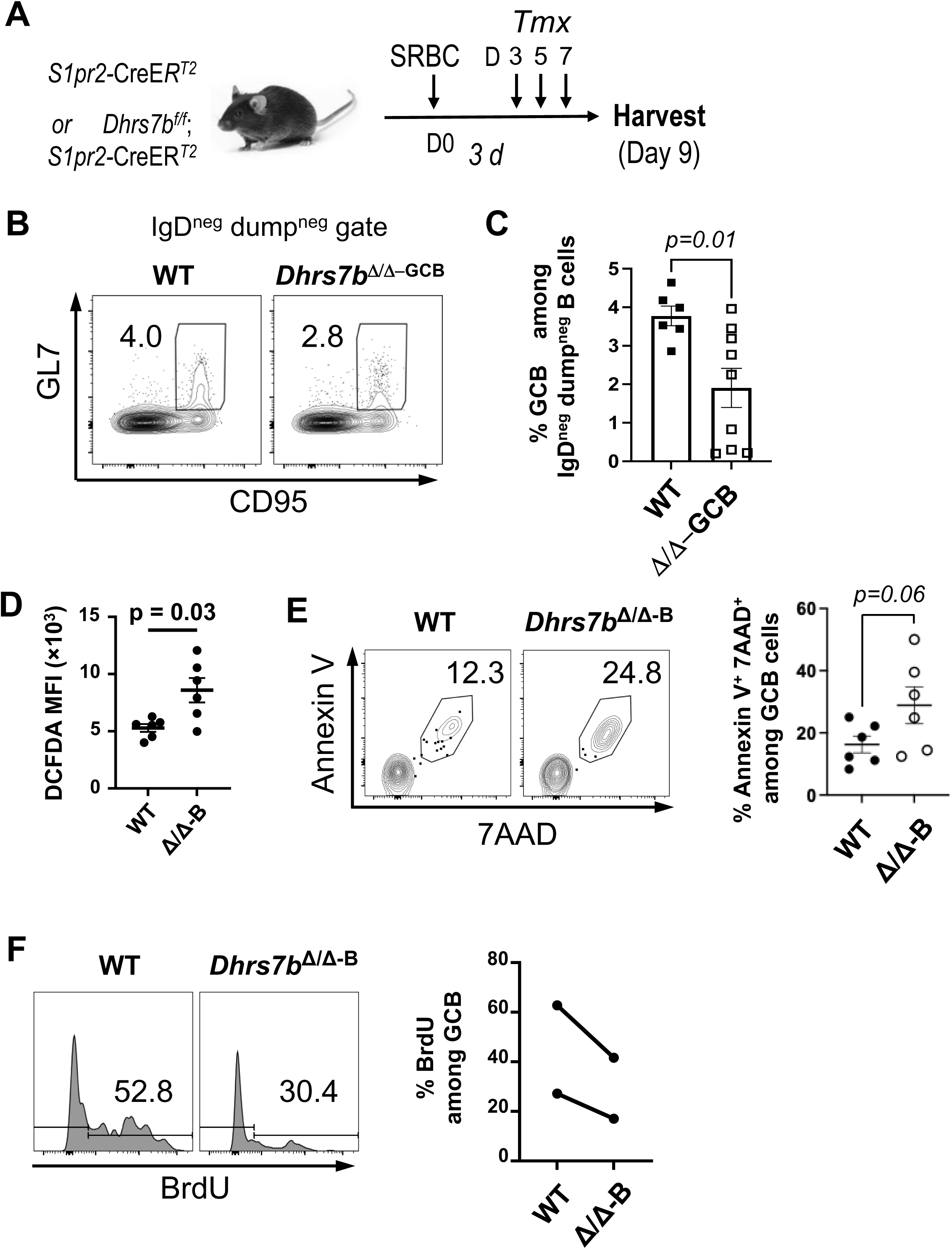
Function of PexRAP in GC response. (A) Schematic of the time line, with immunization followed later by tamoxifen injections into *S1pr2*-CreER^T2^ mice (*Dhrs7b^+/+^*or *Dhrs7b^f/f^*). Note that deletion is only initiated just as the germinal center reaction starts (∼ 3.5 d post-immunization). Mice (*S1pr2*-CreER^T2^; *Dhrs7b^+/+^*or *S1pr2*-CreER^T2^; *Dhrs7b^f/f^*) were immunized with NP-OVA, treated with tamoxifen on days 3, 5, and 7 after NP-OVA immunization (36) and harvested at day 9. (B) Representative flow plots of GL7*^+^* CD95*^+^* GC B cells among viable IgD^neg^ B cells and (C) aggregated mean (±SEM) frequencies of such GC B cells from three replicate experiments (7 WT; 9 cKO mice). Mann-Whitney U test was used to calculate p values. (D-F) Effect of PexRAP on the levels of ROS (D), cell death (E), and proliferation (F) of GC B cells. (D) Total cellular ROS in IgD^neg^ CD38^neg^ GL7^+^ GC B cells were determined by flow cytometry after staining with surface markers and H_2_DCFDA as described in the Methods. The graph shows the mean (±SEM) geometric MFI of H_2_DCFDA from two independent replicate experiments (n = 6 WT and 6 cKO). (E) PexRAP promotes GC B cell survival. Shown are the representative flow plot (left), and a dot graph aggregating all experiments’ outcomes for the frequencies of annexin V^+^ 7AAD^+^ cells in GC B gated cells as in Fig 7D (right panel). (F) PexRAP regulates proliferation of GC B cells. Tamoxifen-treated mice (*huCD20*-CreER^T2^; *Dhrs7b^+/+^*or *huCD20*-CreER^T2^; *Dhrs7b^ff^*, i.e., *Dhrs7b^Δ/Δ-B^*) were immunized with SRBC, and the mice were injected with BrdU as described in Methods. Shown are representative histograms for WT and *Dhrs7b^Δ/Δ-B^* GC B cells as indicated (left panel) and a graph indicating the aggregated result of each independent experiments (right panel) (x=2; n = 6 WT and 6 cKO).

### *Dhrs7b* contributes to modulation of ROS and their impact on B cell population growth

Increased susceptibility to cell death after activation would be a mechanism that could cause the reduced population expansion, GC, and Ab production. Consistent with earlier analyses of LPS-stimulated B cells (70), we had noted that ROS levels were higher in GC B cells, memory B cells (MBCs) and ASCs than in the naive B2 B cell pool (36). ROS can be produced in activated B cells via diverse sources that include mitochondrial electron transport chain function, NADPH oxidase activated by BCR engagement, and fatty acid oxidation in peroxisomes (71–73). ROS can positively mediate signal transduction, but their steady-state concentrations need to be calibrated because overproduction can have deleterious effect on cells, for instance via lipid peroxidation at excessive rates (74–76). Endogenous antioxidant properties have been imputed to plasmalogens because the vinyl ether bond is susceptible to reaction with reactive oxygen, thereby scavenging ROS (48, 57–59), so we tested if *Dhrs7b* function contributes to redox control in B cells. ROS levels were ∼2-fold higher in GC B cells from immunized *Dhrs7b*^Δ/Δ-B^ mice compared to those from immunized WT controls (Fig 7D). *Dhrs7b*^Δ/Δ^ GC B cells also showed higher cell death and attenuated BrdU incorporation (Fig 7E, F; Supplemental Fig 6C.) Moreover, CD40-stimulated *Dhrs7b*^Δ/Δ^ B cells exhibited ∼3 fold higher signal and about a doubling when stained with fluorescent sensors of cellular and mitochondrial ROS, respectively, compared to WT B cells *in vitro* (Fig. 8A-D). Substantially increased ROS also were measured after activation by BCR and CD40 cross-linking (Supplemental Fig 6D). Iron- and copper-dependent lipid peroxidation is considered to be a biological mechanism for ROS-mediated cell death (75, 76). Lipid peroxidation measured by a fluorescent indicator (C11-Bodipy) in vitro was higher in *Dhrs7b*^Δ/Δ^ B cells compared with WT controls after each type of mitogen activation (Fig. 8E; Supplemental Fig 6E). Chemical elicitation of increased ROS and lipid peroxidation in B cells, independent from B cell activation or proliferation, rapidly led to reduced cell viability and B cell numbers (Fig. 8F; Supplemental Fig 7A). Furthermore, in vitro analyses indicated that B cells lacking PexRAP also exhibited increases in the activated executioner caspase, cleaved caspase-3 (CC3) along with early apoptotic cells that are annexin V^+^ but exclude 7-aminoactinomycin D (Fig. 8G, H). These data indicate that *Dhrs7b* function supports not only protection against cell death, but also control of cellular and mitochondrial ROS levels and their impact on lipid modification (Supplemental Fig. 7). Of note, while H_2_O_2_ did not increase this early apoptotic population (Supplemental Fig. 7C), increasing ROS with menadione drove B cell death preceded by annexin V^+^ state without an increase in C11-Bodipy signal (Supplemental Fig. 7D-G). We infer that rather than a single mode of death, increased ROS resulting from PexRAP depletion probably stimulates at least two distinct modes of B cell death.

**Figure 8.**
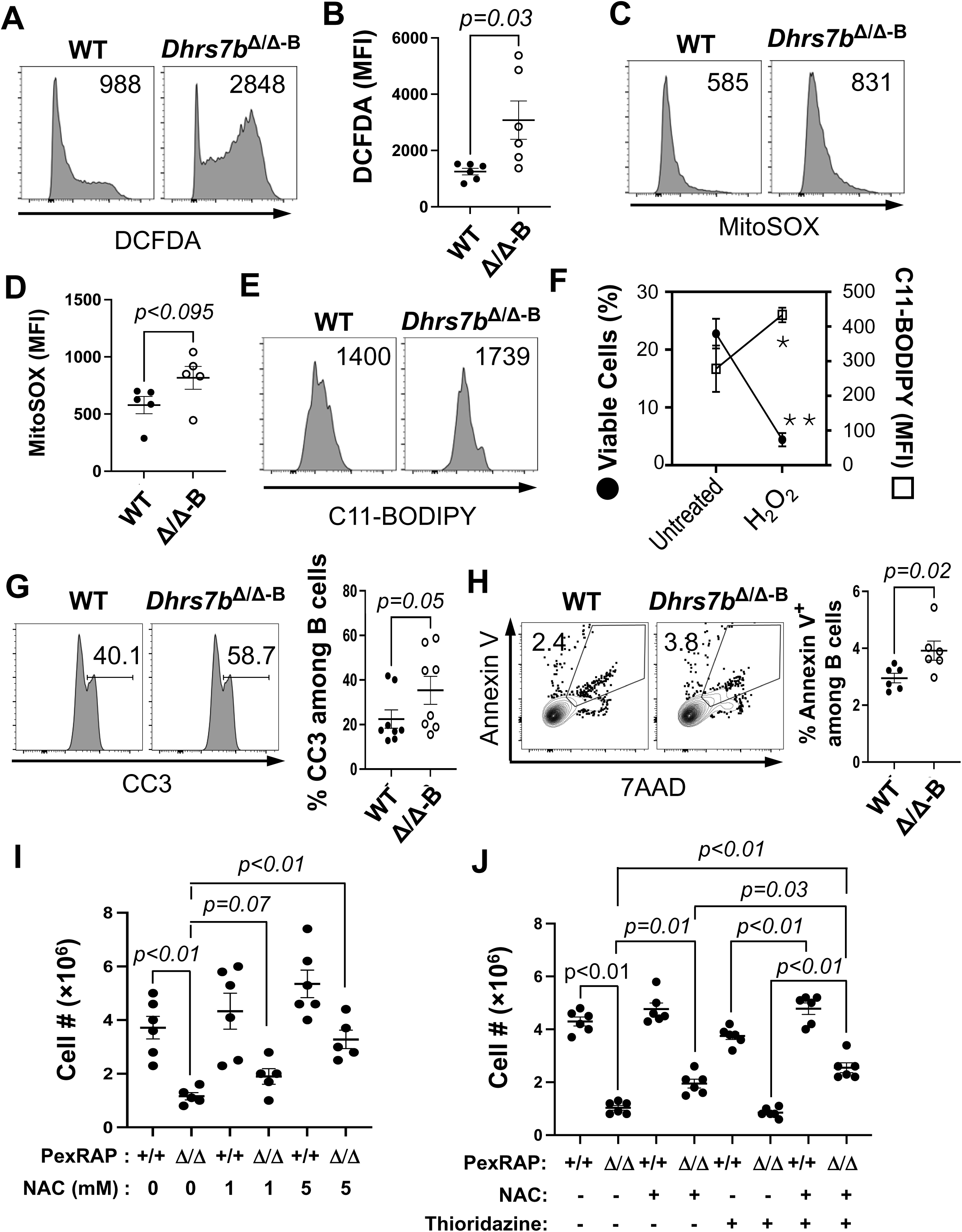
PexRAP contributes to ROS homeostasis and B cell population growth in vitro. (A-D) PexRAP is critical for maintenance of normal ROS levels. Bead-purified B cells from spleens of tamoxifen-treated *huCD20*-CreER^T2^ mice (*Dhrs7b^Δ/Δ-B^* and *Dhrs7b^+/+^*) were cultured 3 days in anti-CD40, BAFF, IL-4, IL-5 and 4-hydroxytamoxifen. Total cellular (A, B) and mitochondrial ROS, mtROS (C, D) in B lymphoblasts were then determined by flow cytometry after staining with surface markers and H_2_DCFDA and MitoSOX, as described in the Methods. Representative histogram image of H_2_DCFDA (A) and MitoSOX (C) in the B cell gate, and aggregated mean (± SEM) geometric MFI of H_2_DCFDA (B) and MitoSOX (D) from 3 independent experiments, each using two mice of each type (6 WT; 6 cKO). P values were calculated by Mann-Whitney U test. (E) PexRAP restrains lipid peroxidation. B cells were activated and cultured as in (A). A representative result of flow cytometric analyses of lipid peroxidation assayed using C11-Bodipy is shown, based on three independent experiments. (F, G) PexRAP promotes B cell survival. (F) Shown are the mean (±SEM) frequencies of total viable ‘events’ (by FSC, SSC, and 7-AAD exclusion) in flow cytometry (filled circles) and MFI of Bodipy-C11 (open squares) after exposure to H_2_O_2_ (200 µM). (*, ** -p=0.03 and 0.003, respectively). (G-I) WT and *Dhrs7b^Δ/Δ^* B cells were cultured (2 d) in anti-CD40, BAFF, IL-4, IL-5 and 4-hydroxytamoxifen. (G) Increased cleaved caspase 3 (CC3) in PexRAP-deficient cells generated in vitro. Cleaved caspase 3 in B cells was detected by intracellular staining and flow cytometry. Shown are a representative pair of histograms for WT and *Dhrs7b^Δ/Δ^* B cells as indicated (left panel) and a dot graph aggregating all experiments’ outcomes (right panel). (H) Frequencies of apoptotic B cells in cultures as in (A-D) were scored for annexin V and 7AAD as described (84). Shown are representative data of flow plots in the lymphocyte gate (left panel) and a dot graph aggregating all experiments’ outcomes (right panel). (I, J) PexRAP and ROS control promote B cell population growth in vitro. WT and *Dhrs7b^Δ/Δ^* B cells were cultured 5 days in anti-CD40, BAFF, IL-4, IL-5 and 4-hydroxytamoxifen in the presence or absence of ROS scavenger NAC (1 mM vs 5 mM) (H). (I) B cells activated as in (A-G) were cultured 5 d in the presence or absence of NAC (1 mM) or thioridazine (100 nM). The bar graphs show the mean (± SEM) recovered cell number from three independent experiments, each with two independent samples of each genotype (WT; *Dhrs7b^Δ/Δ^*). Complementary results of experiments including anti-IgM for BCR cross-linking are shown in Supplemental Figs 2 and 6.

Consistent with the increased steady-state ROS and death, activated *Dhrs7b*^Δ/Δ^ B cells exhibited a defect in B cell population growth *in vivo* and *in vitro* (Fig. 2; Fig. 8A-H). The well-established ROS scavenger, N-acetyl-L-cysteine (NAC), exerted a concentration-dependent effect that improved the survival and population growth of PexRAP-deficient B cells (Fig. 8I). We exploited the sub-maximal rescue by a lower concentration of NAC (1 mM) to explore if peroxisomal oxidative metabolism might contribute to the toxicity observed when B cells are PexRAP-depleted. Although thioridazine on its own provided no increase in the B cell population growth, its combination with 1 mM NAC treatment improved growth of the *Dhrs7b*^Δ/Δ^ B cell population compared to either agent on its own (Fig. 8J). Analyses of B cells activated by BCR crosslinking along with anti-CD40 also found that PexRAP expression in B cells supports their population increase (Supplemental Fig 6F). Moreover, inclusion of NAC in the cultures partially mitigated the inactivation of *Dhrs7b* (Supplemental Fig 6F). Of note, measurements of CTV partitioning identified a modest but statistically significant decrease in division counts of PexRAP-deficient B cells under these conditions, and found that NAC did not increase this component of proliferation (Supplemental Fig 6G, H). These data support the conclusion that sufficient generation of PexRAP-dependent products is crucial for B cell survival. Moreover, this enzyme contributes - directly or indirectly - to a resolution or detoxification of ROS that is critical for population growth of activated B cells.

### *Dhrs7b* in B cells affects their oxidative metabolism and ER mass

Our finding that mitochondrial ROS were increased prompted us to investigate how PexRAP affects physiological functions by measuring respiration in activated B cells. Mitochondrial stress tests with in vitro activated B lymphoblasts measured small but definite decreases in both basal and maximal respiration (oxygen consumption rates, or OCR) (Fig. 9A-C). The altered performance of mitochondria was associated with reductions in calculated ATP generation and proton leak as well as a small decrease in spare respiratory capacity (SRC) (Fig. 9D-F, respectively). In contrast, the rates of glucose-stimulated extracellular acidification - cytosolic reactions and a surrogate that can approximate glycolytic activity - were unaffected (Fig. 9G). Thus, the effects of PexRAP on B cell metabolism are not global, in that altered respiration did not reflect a decrease in glucose-stimulated acidification. Inasmuch as peroxisomes and mitochondria form contacts and functionally interact with the ER, we measured relative ER mass and found that the signal of the fluorophore ERTracker was consistently reduced in PexRAP- depleted B cells (Fig. 9H, I). We infer that both mitochondrial function and ER mass in B cells are promoted by the product of the *Dhrs7b* gene.

**Figure 9.**
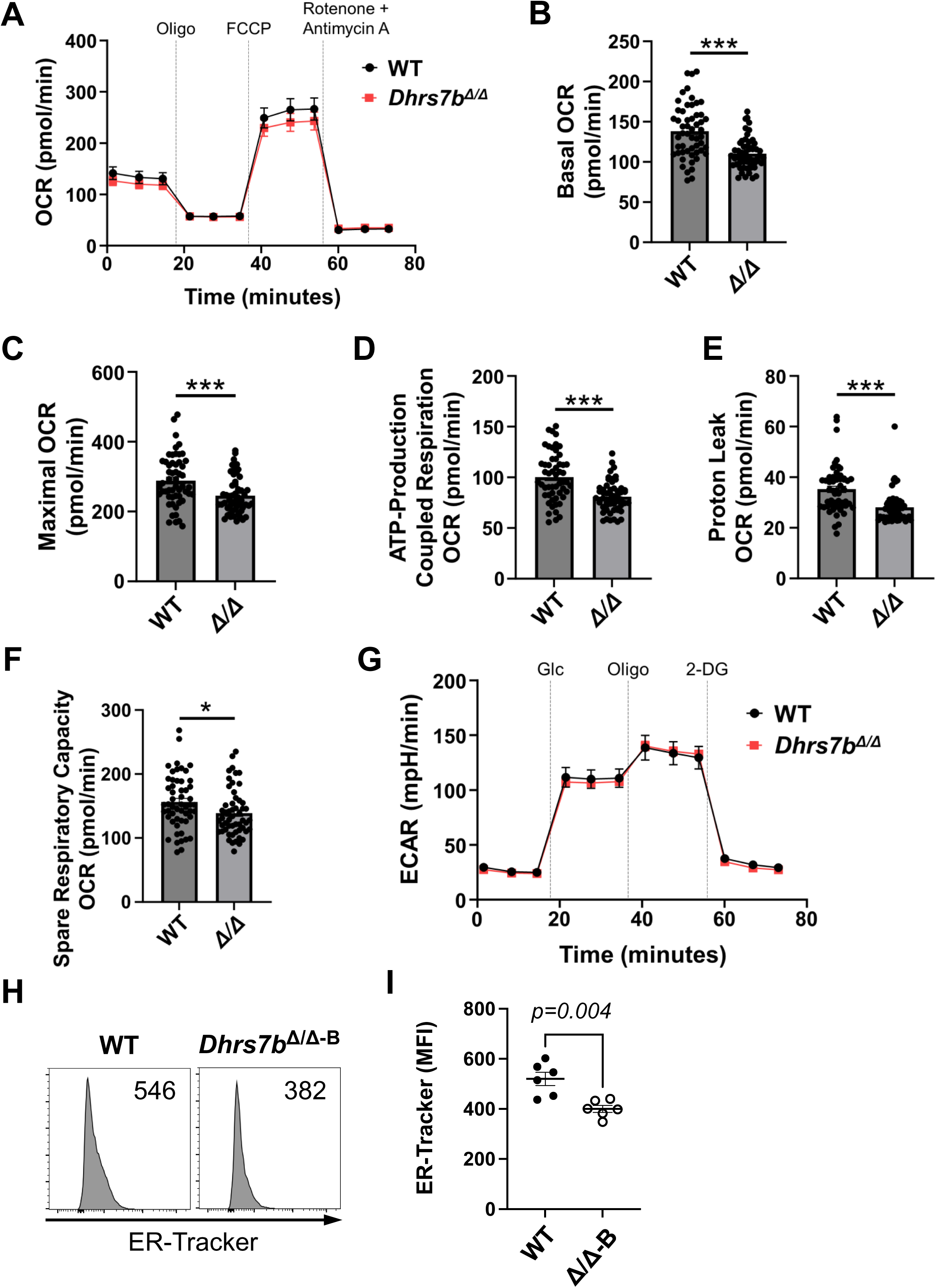
PexRAP deficiency in activated B cells reduces mitochondrial metabolism and ER mass. (A-G) Purified B cells were activated with and cultured (2 d) in anti-CD40, BAFF, IL-4, IL-5, and 4-OHT, then analyzed using a Seahorse XFe96 after harvest and division into equal portions. (A) Shown are aggregated results from three independent experiments measuring oxygen consumption rate (OCR) quantified via metabolic flux analysis during mitochondrial stress testing. (B) Basal OCR, (C) maximal OCR, (D) ATP-production coupled respiration, (E) proton leak, and (F) spare respiratory capacity of WT and *Dhrs7b^Δ/Δ^*B cells assayed in (A). Data points of individual samples are show (each individual dot), as well as bars to display mean (±SEM) values. T-tests with Welch’s correction were used to calculate p-values. *, p<0.05; ***, p<0.001. (G) Extracellular acidification rate (ECAR) of B cells cultured as described previously (36) and assessed using the glycolytic stress test. (H, I) PexRAP influences ER mass. WT and *Dhrs7b^Δ/Δ^* B cells were cultured 5 days in anti-CD40, BAFF, IL-4, IL-5 and 4-hydroxytamoxifen followed by the staining with ER-Tracker Green. Representative histogram of ER-Tracker Green in viable cells (H) and aggregated MFI (±SEM) of ER-Tracker Green from three replicate experiments (I). Mann-Whitney U test was used to calculate p values.

## DISCUSSION

We have shown herein that B lymphocyte expression of an enzyme essential for biosynthesis of a subset of ether phospholipids is crucial for part of the increases in their levels localized to GC. Moreover, the genetic approach used to deplete B cells of PexRAP allowed us to show that the B cell type-restricted gene, *Dhrs7b*, promotes B cell survival, proliferation, and GC size along with affinity maturation and serum concentrations of Ag-specific Ab after immunization. Taken together, this work (i) establishes the predictive value of the discovery approach that identified an unexpected feature of cellular biochemistry in GC, (ii) indicates that B cell-autonomous generation of ether lipid species is crucial for B cell survival, and (iii) regulates the qualities of the Ab elicited by immunization.

Using imaging mass spectrometry for discovery-based hypothesis generation, we had reported that the concentrations of at least a dozen ether phospholipids are increased in splenic GC relative to the remainder of the tissue (45). The analyses presented here confirm and extend the observations, and provide evidence that after immunization, these concentrations increase even in the primary B cell follicle, albeit to a modest degree when compared to the magnitude of increases in GC, and in B cells activated ex vivo. The cell type-specific gene inactivation after establishment of a pre-immune population establishes that an enzyme crucial for generation of a number of plasmalogens has substantial effects on B lymphocyte physiology and function. Plasmalogens and other ether lipids and phospholipids have long been shown to be major constituents of lipid bilayers and cell membranes (48, 49, 57). However, remarkably little is known about whether or not cell-intrinsic synthesis of ether lipids or the balance among their specific molecular species matters for hematopoietic cells or in immunity. Several inborn errors of metabolism in humans are attributable to loss-of-function mutations of genes encoding peroxisomal proteins that impact ether lipid (and hence plasmalogen) synthesis (58) but affect other critical processes as well [reviewed in (48, 58–62)]. Several of these - such as Zellweger Syndrome and rhizomelic chondrodysplasia punctata (RCDP) - and the mouse models generated to study mechanisms in these human disease, lead to severe neurological defects and early post-partum death (57, 58). Whether or not lymphocyte development, homeostasis or function were affected in these studies is not clear. Deficient generation of ether lipids is among many abnormalities caused by elimination of fatty acid synthase (FAS) (46, 47). Acute inactivation of *Fasn*, the gene encoding this enzyme, in young mature mice caused a decrease in spleen size disproportionate to the reduction in neutrophils, which was the most profound hematological consequence (46). In addition, circulating lymphocytes were reduced despite normal steady-state representation in the marrow. In-trans cell-extrinsic functions of ether lipids have been noted previously - specifically, as ligands for the invariant natural killer T cell receptor (77) or for a G-protein coupled receptor on natural killer cells (78). As such, the widespread loss of FAS function and the potential for indirect and pleiotropic effects of this enzyme deficiency left unresolved what cells actually depend on their own synthesis of ether lipids.

PexRAP, the enzyme encoded by the *Dhrs7b* gene, generates intermediates vital for the synthesis of some - but not all - ether lipids in cells such as neutrophils (46). We interpret our lipidomic data as indicating that this is the case for B lymphocytes. Specifically, although a number of plasmalogen or other ether lipid species in the naive, activated, or, in IMS, GC B cells were lower as a consequence of cell type-specific gene inactivation, other phospholipids in these classes were unaffected. Analyses of neutrophil numbers and survival provided evidence that this enzyme can be crucial for a normal membrane composition. Moreover, PexRAP promoted viability of this very short-lived cell type whose membranes have a high plasmalogen content (46). The bloodborne population of lymphocytes was reduced in the *Rosa*26-CreER^T2^, *Dhrs7b*^f/f^ mice (46), but the specific cell types were not reported and cells in circulation may not reflect the spleen or other organs. Of note, the interpretation that the neutrophil phenotype is due to insufficiency of ether lipids or plasmalogens has been questioned based on alternative gene disruption results (54). Several models of unconditional gene inactivation or mutation that drastically reduced the phospholipid end-products were noted to have normal neutrophils [summarized in (54), albeit at older ages (4 - 7 mo) than those used in (46). However, these caveats are subject to points articulated in (53). There may be age-related changes in the processes analyzed herein using young (age ∼ 2 mo) mice (53). Moreover, cellular and organismal selection for adaptation variants makes the results of acute loss-of-function inherently different from unconditional loss and heterozygote parentage (53). The potential for toxicity induced by 4OHT activation of CreER^T2^ was among concerns mooted about the earlier study of neutrophils (54). This concern may have been extrapolated from papers that noted impairment in earlier hematopoiesis when *Rosa26*-CreER^T2^ was combined with very intense regimens of tamoxifen administration to infant mice (65, 79). Such studies used individual doses over 4-fold higher that those used herein, and delivered by gavage five times on a daily basis instead of three. Nonetheless, the IgM^+^ B cell population sizes in spleen and marrow were normal even in the face of reduced erythropoiesis and thymopoiesis (65). Moreover, the analyses herein compared B lineage-specific deletion to CreER^T2+/-^, *Dhrs7b*^+/+^ mice identically dosed in parallel with the test subjects. Thus, our work indicates that PexRAP is essential for normal physiology and function of mature B lymphocytes, in particular after their activation.

The findings are most consistent with a capacity for PexRAP function to contribute towards B lymphocyte survival and effective proliferation by restraining both executioner caspase activation and lipid peroxidation due to excessive ROS levels. For instance, population expansion of *Dhrs7b* Δ/Δ B cells was severely undermined after mitogenic stimulation in vitro with little difference in the division history but higher frequencies of annexin V-positive B cells. These findings are consistent with those obtained with neutrophils (46). On the surface such results might appear to present a conundrum relative to work indicating that both fatty acyl-CoA reductase 1 and the enzyme that later generates an alkenyl bond at a terminal step in plasmalogen biosynthesis are crucial for the susceptibility of HT-1080 and 786-O tumor cell lines to ferroptosis (55). However, our data are consistent with published evidence that PexRAP is needed for a narrower subset of ether lipids than the broad requirement for FAS or fatty acyl-CoA reductases (46–49, 80, 81). Moreover, molecular dissection of mechanisms in various non-hematopoietic cell types (including the cancer line 786-O) showed that ether lipid biosynthesis can either promote susceptibility or resistance to ferroptosis (82). Collectively, our findings suggest that for B cells the inactivation of PexRAP tilts the balance toward death and reduced Ab - perhaps involving lower fractions of poly-unsaturated fatty acids (PUFA) (82), or a lower ratio of plasmalogens (alkenyl linkages) relative to alkyl-linked ether lipids.

PexRAP localizes to peroxisomes (35–37), and prior work has suggested both that peroxisomes are increased in GC B cells and likely contribute to FAO by these cells (26). In line with the possibility that this organelle may affect PexRAP-dependent survival, our data indicated that specific inhibition of peroxisomal FAO could collaborate with sub-optimal ROS scavenging to mitigate the reduced growth of activated B cells. However, recent findings indicate that PexRAP can also be expressed in the ER and is capable of contributing to ether lipid biosynthesis via catalysis in this organelle in addition to or instead of the peroxisome (56). Thus, although ROS generated in peroxisomes likely contribute to the stress experienced by PexRAP-deficient B cells, the phenotypes observed herein may not exclusively reflect a need for the peroxisomal fraction of the enzyme but instead might include functions of ER-localized protein.

Our findings with an intermediary in ether lipid biosynthesis add to an increasing body of evidence that metabolism in B lymphocytes affects their differentiation or function (28–36). Many sources and forms of Ab can be protective, while in several autoimmune diseases these molecules can drive pathology (1, 2). The capacity to generate increased affinities and circulating concentrations of Ab is an important factor both in protective immunity and auto-immune disease. Antigen-independent, TLR-driven PC differentiation requires increased oxidative phosphorylation (70), while oxidative metabolism in GC B cells promotes affinity maturation by unknown mechanisms (33–35). While not the exclusive determinant of such properties, GC are major sources of higher affinity Ab with a broader spectrum of specificities (9, 13, 14). A limitation of our study is that for practical reasons, we were unable to use purified GC B cells for immunoblots or nucleic acid quantitation to determine the extent of counterselection or extent of Dhrs7b locus inactivation in these cells after their clonal expansion. Nonetheless, several lines of evidence suggest that PexRAP affects the physiology of GC B cells during their residence in this micro-anatomic structure. Ether lipids are considered to be short-lived, and the levels of several within the GC were reduced by PexRAP depletion prior to immunization. Moreover, rates of cell cycling as measured with BrdU incorporation appeared reduced, while ROS levels within GC B cells were higher along with the frequency of death. Because GC Tfh cells provide new stimulation to GC B cells that present peptide recognized by the TCR, some of these effects may be secondary consequences of a substantially lower GC-Tfh population even when B cell-specific loss-of-function was tested. A speculative corollary is that the observed reduction in ER mass might be functionally notable. For instance, cognate B-T interaction in the GC involves BCR-mediated internalization of Ag followed by processing and MHC-II presentation of T cell epitope peptide. Moreover, ether lipids and plasmalogens are presented to innate-like T cells via CD1d (77). In addition to the GC-localized abnormalities, GC formation, size, and function also depend on the efficiency of population growth for B cells in a phase before they enter and take on characteristics of GC B cells, and then homeostatic and differentiative processes while in the secondary follicle. Thus, while the findings with the post-immunization inactivation of *Dhrs7b* using *S1pr2*-CreER^T2^ provide evidence that the expression of PexRAP is important within GC B cells, the decreases of GC and affinity-matured Ab are likely also to involve a pre-GC phase of clonal expansion. In any case, the rapidity of the effects suggests that pharmacological inhibition of PexRAP may be a target in inflammatory disease in which B cells participate, particularly in light of the potential for impacts on pro-inflammatory neutrophils.

## MATERIALS & METHODS

### Reagents

Monoclonal antibodies (mAb) against mouse CD40, CD90.2, B220 (CD45R), and other mAbs (purified, biotinylated, or fluorophore-conjugated) were from BD Biosciences or Tonbo Biosciences (San Diego CA) unless otherwise indicated. BAFF was from AdipoGen (San Diego, CA). Recombinant mouse IL-4 and recombinant mouse IL-5 were from Peprotech (Rocky Hill NJ). Glucose, 2-deoxyglucose, oligomycin, rotenone, antimycin A, carbonyl cyanide 4-phenylhydrazone (FCCP), N-acetyl-cysteine (NAC), thioridazine hydrochloride, H_2_O_2_, and 4-hydroxytamoxifen (4-OHT) were from Sigma-Aldrich Chemicals (St. Louis MO), and menadione from Cayman Chemicals (Ann Arbor, MI). Tamoxifen (Tmx) was from APExBio Technology (Houston TX). NP-BSA and NP-PSA (for capture in ELISA) as well as NP-OVA (for immunization) were obtained from Biosearch Technology (Novato CA). SRBCs (sheep red blood cells) were from Thermo Fisher Scientific (Waltham MA). CellTrace^TM^ Violet cell proliferation kit, CM-H_2_DCFDA, MitoSOX^TM^ Red, and BODIPY^TM^ 581/591 C11 were obtained from Invitrogen (Waltham, MA).

### Mice, immunizations, ELISA and ELISpot

All animal protocols were reviewed and approved by the Vanderbilt University Institutional Animal Care and Use Committee. Mice were housed in ventilated micro-isolators under Specified Pathogen-Free conditions in a Vanderbilt University Medical Center mouse facility and used both male and female mice at 6–8 weeks of age. For cell type specific inactivation of *Dhrs7b* genes, *Dhrs7b*^f/f^ mice (46) were crossed with *Rosa26-*CreER^T2^ (46), *huCD20-*CerER^T2^ (30), or *S1pr2-*CreER^T2^ strains (51, 52), all on C57/BL6 genetic background. Tamoxifen was administered as previously reported (25, 30). To control for potential Cre toxicity, CreER^T2^ mice were similarly injected to use as wild-type (WT) controls. Mice (ages ∼ 8wk) were immunized with SRBCs (2×10^8^ cells per mouse) and analyzed 1 week after immunization as described (25, 45). Alternatively, mice were immunized and boosted by i.p. injections [each of 100 µg NP_16_-OVA (Biosearch Technologies, Novato, CA) emulsified in 100 µL of Imject^TM^ Alum (Thermo Fisher Scientific, Pittsburgh, PA)] as described previously (25, 30), and harvested for analyses 1 wk after the 2^nd^ injection. Isotype-specific relative levels of Ag-specific Ab were quantitated by capture ELISA using Ag (NP_20_-BSA or NP_2_-PSA for all- or high-affinity hapten-specific Abs, respectively) followed by SBA Clonotyping System (Southern Biotech, Birmingham AL), as described previously (25, 30). As previously described (30), frequencies of Ab-secreting cells (ASCs) were analyzed by ELISpot and quantitated using an ImmunoSpot Analyzer (Cellular Technology, Shaker Heights OH).

### Cell cultures, proliferation assay, and reversion of population growth

Splenic B cells were purified (90-95%) using negative selection as previously described (30) or by positive selection with anti-mouse B220 microbeads (Miltenyi Biotech, Auburn, CA). B cells (5 × 10^5^ cells in 1 ml) were activated with anti-CD40 (1 μg/mL) and cultured with BAFF (10 ng/mL), IL-4 (10 ng/mL), IL-5 (10 ng/mL) and 4-OHT (50 nM) in Iscoves Modified Dulbecco’s Medium (IMDM) supplemented with 10% Fetal Bovine Serum (FBS), 100 U/mL penicillin 100 µg/mL streptomycin (Invitrogen), 3 mM L-glutamine, nonessential amino acids (Invitrogen), 0.1 mM 2-ME (Sigma-Aldrich). To analyze the effect of ROS scavenger and inhibition of peroxisomal lipid oxidation on population growth, bead-purified B cells from tamoxifen-treated *huCD20-*CerER^T2^ or *Dhrs7b^f/f^*; *huCD20-*CerER^T2^ mice were activated with anti-CD40, cultured with BAFF, IL-4 and IL-5 in the presence or absence of NAC (1 mM and 5 mM) and/or thioridazine·HCl (100 µM) for 5 days. For enhancement of ROS in WT B cells, purified B cells were analyzed 3 d after activation as for ROS scavenging, but vehicle, H_2_O_2_ (200 μM) or menadione (8 μM) was added at three different time points (overnight, 3 hours, and 1 hour prior to processing for flow cytometry to counting and measurements of DCFDA, annexin V, and C11 Bodipy signals.

For the analysis of proliferation in vitro, B cells were purified by depleting CD90.2^+^ T cells and CD138^+^ cells using biotinylated anti-CD90.2 Ab and biotinylated anti-CD138 Ab followed by streptavidin-conjugated microbeads (iMag^TM^; BD Bioscience), labeled with 5 µM CellTrace^TM^ Violet (CTV, Invitrogen), activated and cultured as above. To analyze proliferation in vivo, B cells were purified, labeled with CTV, and injected intravenously into B cell deficient µMT recipient mice. Mice were harvested at 4 days after adoptive transfer. To measure proliferation rates of GC B cells, in vivo BrdU incorporation was performed by injecting mice with BrdU (2 mg/mouse; BD-Pharmingen, San Jose, CA) at 16 h and 4 h before harvesting spleens from SRBC-immunized mice. Cells were stained for surface markers (B220, IgD, GL7, CD38) followed by fixation, permeabilization, and staining with anti-BrdU-APC Ab using the BrdU Flow Kit (BD-Pharmingen) according to manufacturer’s protocol.

### Flow cytometry - general and measurements of ROS, lipids peroxidation, and ER

GC B cells were identified GL7^+^ CD95^+^ events in the viable B220^+^ dump^-^ gate (dump channel consisting of one fluorophore for CD11b, CD11c, F4/80, Gr1, TCRβ, and 7AAD), while plasmablasts were defined as B220^+^ CD138^+^ TACI+ dump^-^ cells. For flow analyses of total intracellular ROS and mitochondrial superoxide, B cells (1 × 10^6^ cells) were washed with PBS and stained with 1.25 µM CM-H_2_DCFDA or 5 µM MitoSOX^TM^ Red in PBS (20 min at 37°C), respectively, then washed with 1% BSA containing PBS, and further stained with anti-B220, anti-CD19, anti-CD138, and Ghost-BV510. To measure the lipid peroxidation, cultured B cells (1 × 10^6^) were washed with PBS and stained with 1.25 µM C11-BODIPY^TM^ in PBS (20 min at 37°C), and washed with 1% BSA containing PBS followed by surface staining as above. To analyze the level of endoplasmic reticulum (ER), B cells (1 × 10^6^) were stained with 0.5 µM ER-Tracker^TM^ Green (Molecular Probes, Eugene OR) in Hank’s Balanced Salt Solution with calcium and magnesium (HBSS/Ca/Mg; Gibco) for 30 min at 37°C.

For measurements of cell death (78), cultured B cells (1 x 10^6^) were washed twice with PBS, once with Annexin V binding buffer (10 mM HEPES, pH 7.4, 140 mM NaCl, 2.5 mM CaCl_2_), and then incubated (15 min at 20 C in the dark) with Annexin V-rPE, 7AAD, APC-conjugated CD138, APC-Cy7-conjugated B220, e450-CD19. Cells were then washed with Annexin V binding buffer and analyzed on the flow cytometer. To measure levels of cleaved caspase-3 (25), an activated apoptosis executor, the cultured cells were rinsed with PBS after harvest, stained (15 min at 4 C) with 7AAD, APC-conjugated CD138, APC-Cy7-conjugated B220, e450-CD19, washed (FBS, 1% v/v in PBS), then fixed with paraformaldehyde (4% w/v, 10 min at 20 C), permeabilized with permeabilization buffer (saponin, 0.2% w/v with FBS 1% in PBS), stained with FITC-conjugated Ab specific for cleaved caspase-3 (BD Biosciences), rinsed, and analyzed by flow cytometry.

### Immunohistochemistry, MALDI-IMS, and IMS data analysis

Mice were immunized with SRBC and spleens were harvested at 1 week after immunization. Spleens were snap frozen on dry ice, and mounted to a cryostat chuck with a minimal amount of OCT (Thermo Fisher Scientific, San Jose, CA). Frozen spleens were cut at 12 µm thickness using a Leica CM3050S (Leica, IL, USA) at -20 °C, fixed with 1% paraformaldehyde for 10 min, washed twice with PBS, and blocked with M.O.M^TM^ (Vector Lab) followed by incubation with GL7-FITC, anti-IgD-PE, and anti-CD35-biotin Ab followed by streptavidin-conjugated Alex647at 4 °C. After the tile scanning of spleen sections, GC size and numbers were quantified with FIJI Image J software.

For Matrix-Assisted Laser Desorption/Ionization (MALDI) Imaging Mass Spectrometry (IMS), frozen spleens were cut as above, and mounting onto indium tin oxide (ITO) coated microscope slides (Delta Technologies ETC). Slides were then vacuum desiccated for at least 30 minutes before matrix application. Pre-IMS autofluorescence (AF) images were acquired prior to matrix application on a Zeiss Axio Scan Z1 Slide Scanner using the brightfield channel and the DAPI (ex. 340-380 nm; em. 435-485 nm, blue), EGFP (ex. 450-490 nm; em. 500-550 nm, green), DsRed (ex. 538-562 nm; em. 570-640 nm, red) filter cubes. After taking AF images, a custom in-house developed sublimation device was used to apply the matrices 2,5-dihydroxyacetophenone (DHA) and 1,5-diaminonaphthalene (DAN) (Sigma Aldrich, St. Louis, MO, USA) to tissue sections for positive and negative ion mode analyses, respectively (78). For immunized AID-GFP mice, the fluorescence emissions identified GC localization within the section used next for IMS. Alternatively, two serial sections of immunized mice spleens were used, one for 2D-IMS and the contiguous section for immunohistochemistry (IHC). Fluorescence data were registered with the 2D-IMS visualizations of phospholipids to align images (45), allowing quantitation of the ion intensities in specific m/z peaks within defined micro-anatomic portions of the spleen (Fig. 4B). MALDI IMS data were acquired with a 10 µm pixel size (laser spot size 8 µm) in full scan mode using a Bruker trapped ion mobility time-of-flight (timsTOF) Flex mass spectrometer (Bruker Daltonics Billerica, MA, USA). Data were acquired with 250 shots per pixel and a mass range of *m/z* 200-2000. SCiLS (software Bruker Daltonics) was used to process the data, normalize ion intensity, visualize ion images, and merge the images.

### LC-MS for lipidomics

To prepare samples, a one-phase method was used to extract lipids (85). Briefly, 0.5 mL of MeOH/MTBE/CHCl_3_ mix (1.3:1:1) was added to a frozen pellet of B-cells (1x10^6^ cells total), spiked with 10 uL EquiSPLASH-lipidomics internal standard mix (Avanti Research), briefly vortexed and shaken gently for 20 min, followed by centrifugation at 20,000 x *g* for 15 min at 10°C. The supernatant was transferred to a clean Eppendorf tube, evaporated under a gentle stream of N_2_ gas, and resuspended in 100 uL methanol/CHCl_3_ (9:1) and 2uL were used for LC-HRMS (high-resolution MS) analysis. Each sample was injected two times - one injection in positive ESI mode followed by one in negative mode. Pooled QCs were injected to assess the performance of the LC and MS instruments at the beginning, in the middle and at the end of each sequence. Discovery lipidomics data were acquired using a Vanquish UHPLC (ultrahigh performance liquid chromatography) system interfaced to a Q Exactive HF quadrupole/orbitrap mass spectrometer (Thermo Fisher Scientific).

Chromatographic separation was performed with a reverse-phase Acquity BEH C18 column (1.7 mm, 2.1x150mm, Waters, Milford, MA) at a flow rate of 250 ul/min. Mobile phases were made up of 10 mM ammonium formate and 0.1% formic acid in (A) H_2_O/CH_3_CN (40:60) and in (B) CH_3_CN/ iPrOH (10:90). Gradient conditions were as follows: 0–1 min, B = 20 %; 1–8 min, B = 20–100 %; 8–10 min, B = 100 %; 10–10.5 min, B = 100–20 %; 10.5–15 min, B = 20%. The total chromatographic run time was 15 min. Mass spectra were acquired over a precursor ion scan range of m/z 200 to 1,600 at a resolving power of 60,000 using the following HESI-II source parameters: spray voltage 4 kV (3 kV in negative mode); capillary temperature 250°C; S-lens RF level 60 V; N_2_ sheath gas 40; N_2_ auxiliary gas 10; auxiliary gas temperature 350°C. MS/MS spectra were acquired for the top-seven most abundant precursor ions with an MS/MS AGC target of 1e5, a maximum MS/MS injection time of 100 ms, and a normalized collision energy of 15, 30, 40. High resolution mass spectrometry data were processed with MS-DIAL version 4.70 in lipidomics mode (86). MS1, and MS2 tolerances were set to 0.01 and 0.025 Da respectively. Minimum peak height was set to 30,000 to decrease the number of false positive hits. Peaks were aligned on a quality control (QC) reference file with RT tolerance of 0.1 min and mass tolerance of 0.015 Da. Default lipid library was used (Msp20210527163602_converted.lbm2), solvent type was set to HCOONH_4_ to match the solvent used for separation, and the identification score cut off was set to 80%. All lipid classes were made available for the search. After lipid identification was completed, MS-DIAL results were exported into Excel and cleaned using minimum RSD for QC samples set to 20% and minimum ratio of QC to Blank set to 10. For species identified in both PexRAP-depleted (*Dhrs7b* Δ/Δ) B cells and controls, mean levels (areas under peak curves) and their variance were analyzed in Excel.

### Metabolic flus analyses

Oxygen Consumption Rate (OCR) were measured using Seahorse XF96 extracellular flux analyzer (Agilent Technology, Santa Clara, CA) as described previously (30). Briefly, in vitro activated B cells were washed twice, resuspended in XF Base Media (Agilent Technologies) supplemented with 2 mM L-glutamine, and equal numbers of B cells (2 × 10^5^) were plated on extracellular flux assay plates (Agilent Technologies) coated with 2.5 µg/mL CellTak (Corning) according to the manufacturer’s protocol. Before extracellular flux analysis, B cells were rested (25 minutes at 37°C, atmospheric CO_2_) in XF Base Media. OCR and ECAR were measured before and after the sequential addition of 1.5 µM oligomycin, 0.5 µM FCCP and 0.5 µM rotenone/antimycin A.

## ACKNOWLEDGEMENTS

The experimental work was funded by NIH Grants to VUMC [R01 AI113292 (M.R.B), R01 HL106812 (M.R.B.), followed by R21 AI164760 and R01 AI149722 (M.R.B.) and Vanderbilt University Medical Center Pathology-Microbiology-Immunology departmental funds]. Mass spectrometry imaging was supported in part by NIH grant P41 GM103391 to Vanderbilt University (R.M.C.). Additional support for M. A. J. was provided by the National Science Foundation, NSF DGE-1445197. NIH Shared Instrumentation Grant 1 S10 OD018015 as well as scholarships via the Cancer Center Support Grant (CA068485) and Diabetes Research Center (DK0205930) helped defray costs of Vanderbilt Cores. We thank L. Clark for a critical reading and comments on the revised manuscript, J. Cyster for generously expediting shipment of *S1pr2*-CreERT2 breeding stock, and Vanderbilt institutional cores (High-Throughput Screening; Flow Cytometry Shared Resource; Mass Spectrometry Shared Resource; Cell & Developmental Biology) for equipment, expertise, and assistance.

## Abbreviations used

Ag: antigen
Ab: antibody
WT: wildtype
KO: knockout
MZB: marginal zone B cell
FoB: follicular B cell
PC: plasma cell
Ig: immunoglobulin
GC: germinal center
ETC: electron transport chain
ROS: reactive oxygen species
mtROS: mitochondrial ROS
FAO: fatty acid oxidation
EL: ether lipid(s)
2D: 2-dimensional
MALDI: matrix-assisted laser desorption/ionization
IMS: imaging mass spectrometry
ER: Endoplasmic Reticulum
PexRAP: Peroxisomal Reductase Activating PPARγ
DHAP: dihydroxyacetonephosphate
G3P: glycerol-3-phosphate
CTV: CellTrace Violet
SRBC: sheep red blood corpuscule
TCR: T cell receptor
CD: Cluster of differentiation
AID: Activation-induced deaminase
GFP: Green fluorescent protein
NP-OVA: 3-Nitrophenylacetyl (NP)-ovalbumin (OVA)
ASCs: Ab-secreting cells
CSR: class-switch recombination
MBC: memory B cell
NADPH: nicotinamide adenine dinucleotide phosphate
NAC: N-acetyl-L-cysteine
OCR: oxygen consumption rate
ECAR: extracellular acidification rate
RCDP: rhizomelic chondrodysplasia punctata
FAS: fatty acid synthase
Tmx: tamoxifen
4OHT: 4-hydroxytamoxifen
PUFA: poly-unsaturated fatty acids
2-DG: 2-deoxyglucose
FCCP: carbonyl cyanide 4-phenylhydrazone
BSA: bovine serum albumin
PSA: porcine serum albumin
H2DCFDA: dichlorodihydrofluorescein diacetate
ELISA: enzyme-linked immunosorbent assay
ELISpot: enzyme-linked immunosorbent spot
DHA: dihydroxyacetophenone
DAN: 1,5-diaminonaphthalene
timsTOF: trapped ion mobility spectrometry time-of-flight
SDS-PAGE: sodium dodecyl-sulfate polyacrylamide gel electrophoresis
SEM: standard error of means.

## SUPPLEMENTAL INFORMATION

**Supplemental Figure 1.**
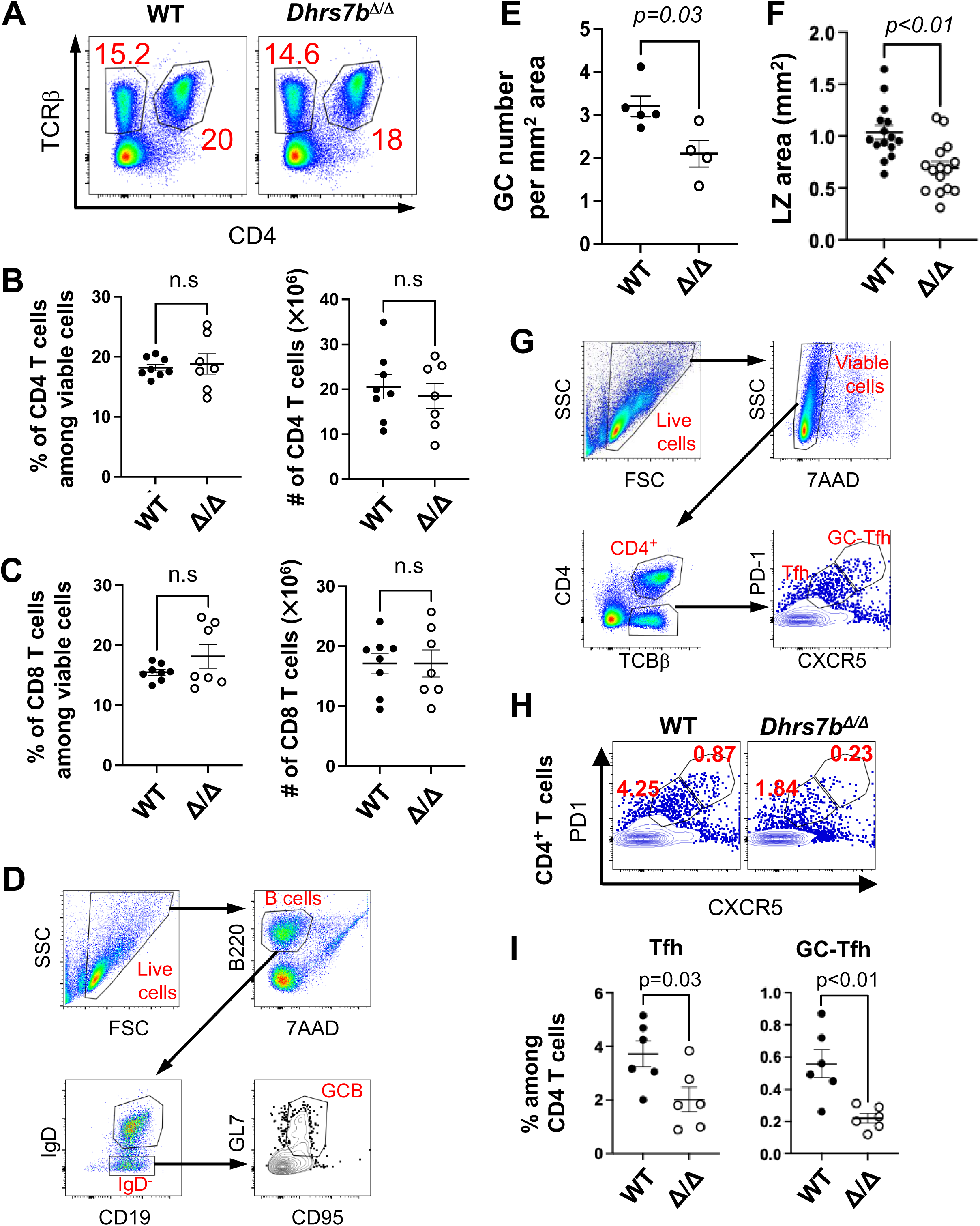
PexRAP is dispensable for T cell numbers, but impacts the Tfh cell population prevalence. Results from flow cytometry analyses of splenocytes from mice (*Rosa26*-CreER^T2^, *Dhrs7b^+/+^,* “WT” or *Rosa26*-CreER^T2^; *Dhrs7b*^f/f^, “*Dhrs7b* Δ/Δ”) harvested after sequential tamoxifen treatments followed by immunization with SRBC and harvest 1 wk thereafter, all as described in *Materials and Methods* and illustrated in Fig. 1A. (A-C) Normal numbers of T cells. Shown are representative flow plots of splenic T cells (A), mean (± SEM) frequencies of TCRβ^+^ CD4^+^ T cells (B), and CD8 (TCRβ^+^ CD4^-)^ T cells (C) from WT and *Dhrs7b* Δ/Δ (cKO) mice based on three replicate experiments totaling 8 WT and 7 cKO mice. (D) Linking to Fig. 1C-F, representative gating scheme illustrating the flow-cytometric determination of splenic B cell frequencies and humbers (overall and GC B phenotype) in mice immunized as in A-C, Fig. 1A. (E, F) Linking to Fig. 1G-I, frequencies of GC, plotted as GC/mm^2^ (E) and sizes of their LZ (plotted as area of each GC in the microscopic sections) (F) identified by immunofluorescent staining and microscopy of spleen sections from immunized mice. (G) Representative gating scheme illustrating the flow cytometric determination of Tfh and GC-Tfh cells in splenocyte suspensions of mice prepared and immunized as in Fig. 1. (H, I) PexRAP promotes Tfh cell population size. (H) Shown are representative flow plots of Tfh and GC-Tfh cells, determined using the gating scheme of Supplemental Fig. 1G. (I) Shown are mean (±SEM) frequencies of PD-1^med^ CXCR5^med^ Tfh cells (left) and PD-1^hi^ CXCR5^hi^ GC-Tfh cells (right) from WT and cKO mice in two independent replicate experiments (n = 6 WT and 6 cKO).

**Supplemental Figure 2.**
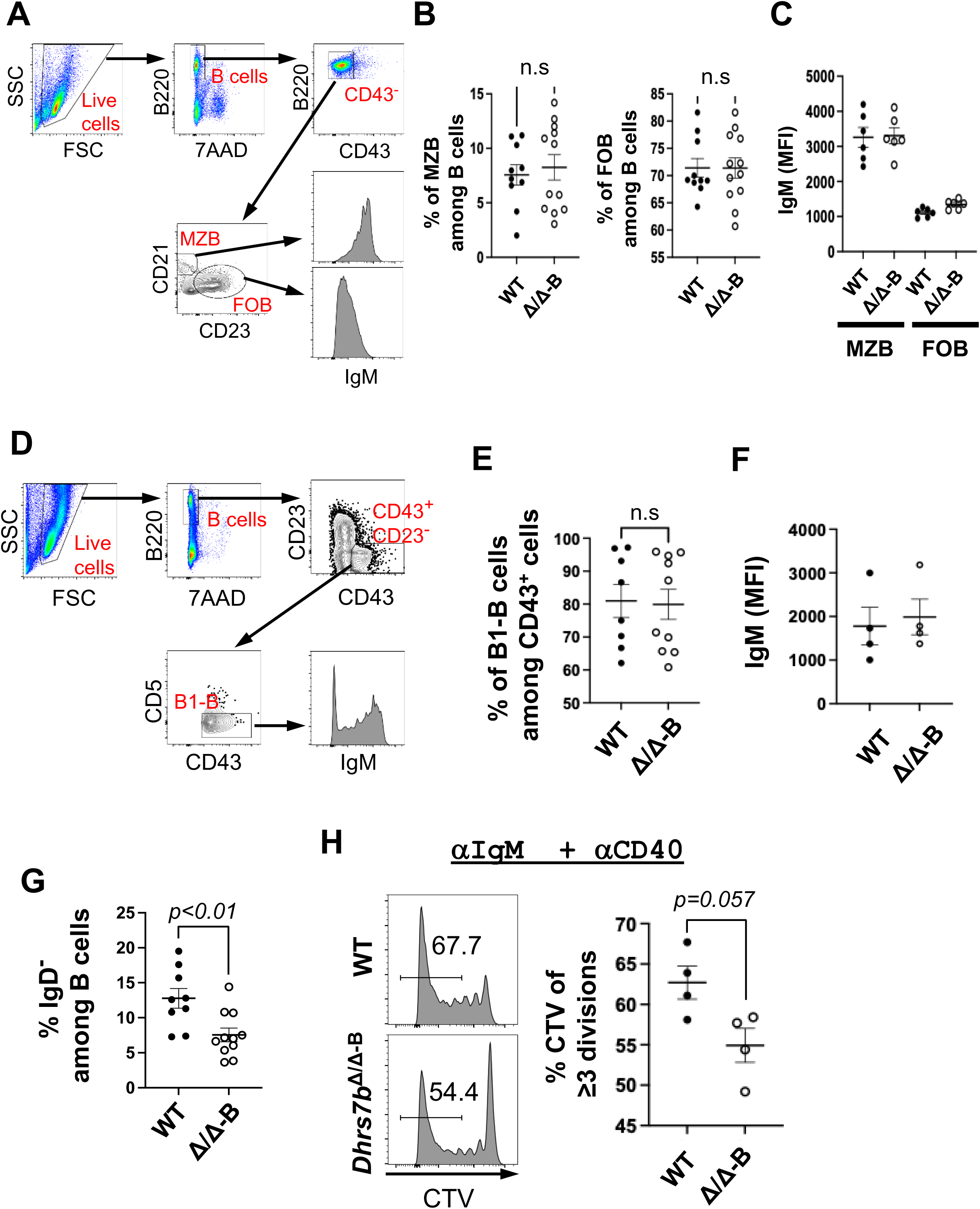
PexRAP is dispensable for pre-immune B cell subset balance and surface IgM expression, but contributes to B cell population growth. (A) A representative gating scheme illustrating the flow-cytometric estimation of the MZ and FO B cell populations in mice. (B, C) Shown are the mean (±SEM) frequencies of B220^+^ CD43^neg^ CD23^neg^ CD21^+^ MZB cells (left panel) and B220^+^ CD43^neg^ CD23^+^ CD21^lo^ FOB cells (right panel) in spleen (B), and aggregated mean (±SEM) geometric MFI of surface IgM on MZB and FOB subsets (C). (D) Gating strategy for identification of B1-b cells and measurement of their surface IgM levels. (E, F) Shown are the mean (±SEM) frequencies of B220^+^ CD23^lo^ CD43^+^ CD5^neg^ B1-b cells in peritoneal cavity (E), and aggregated mean (±SEM) geometric MFI of surface IgM on B1-b cells (F). (G) In vivo generation of IgD^neg^ progeny from transferred B cells of the indicated genotypes, as measured in four independent replicate experiments (n =8 WT and 9 cKO). Shown are mean (±SEM) frequencies of IgD^neg^ B cells as aggregate data for the frequencies of IgD^neg^ events in the gate for viable B cells recovered from µMT recipient mice after transfer of WT or PexRAP-depleted B cells (experiments of Fig. 2D-F), as indicated, with representative flow plots in Fig. 2E. (H) PexRAP promotes B cell proliferation in vitro. B cells were purified and stained with CTV as in Fig 2G. WT and *Dhrs7b^Δ/Δ^* B cells were stained with Cell Trace Violet (CTV), activated and cultured 4 days in anti-IgM, anti-CD40, BAFF, IL-4, IL-5, and 4-OHT. Shown are the representative flow-cytometric analysis of CTV partitioning (left), and aggregated frequencies of divided B cells from two independent replicate experiment, each using two separately sourced pools of B cells (n = 4 WT and 4 cKO). P values were calculated by Mann-Whitney U test.

**Supplemental Figure 3.**
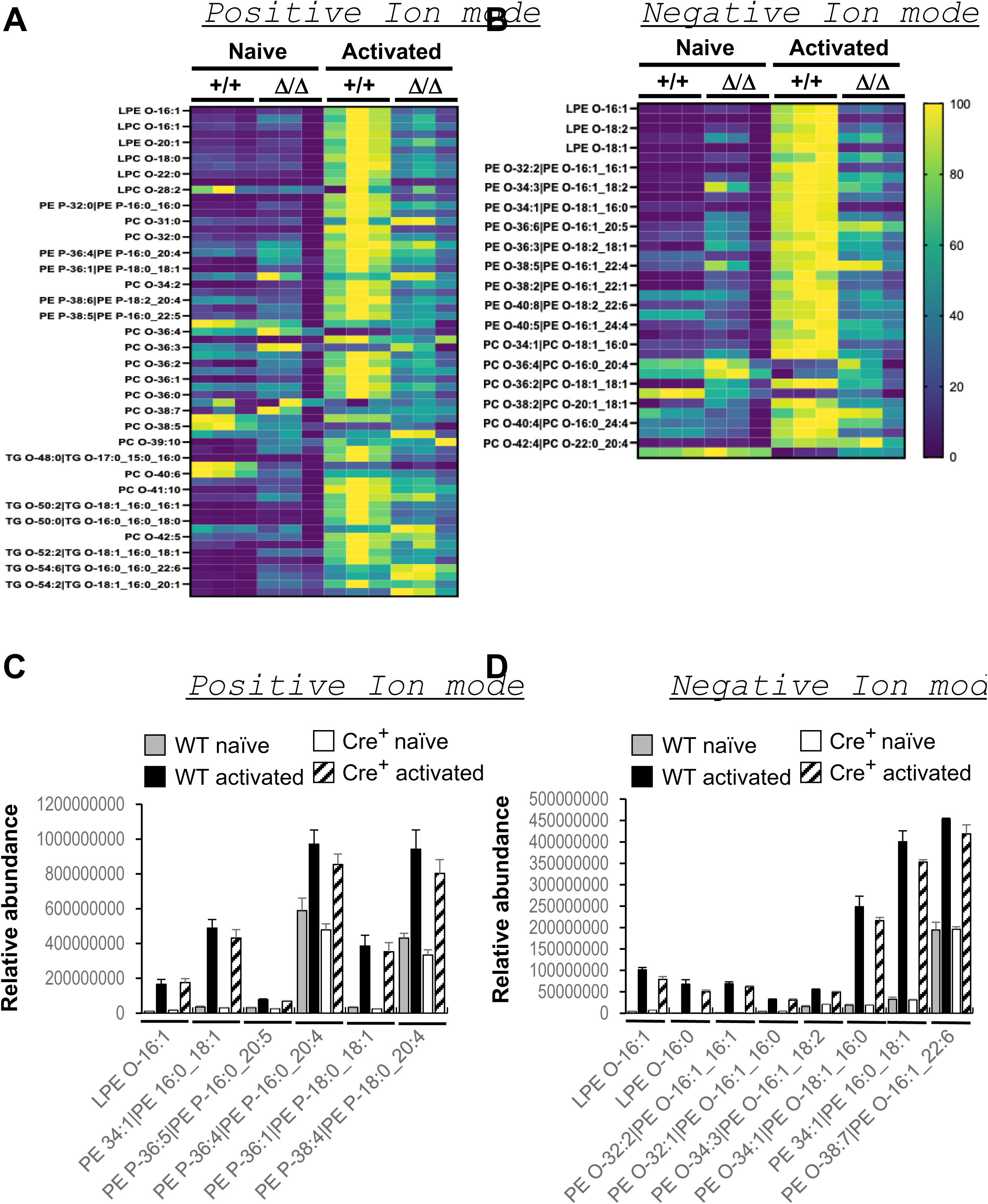
Activation of B cells induces PexRAP-dependent increases in their ether and plasmalogen P-lipids. As in Fig. 3, B lymphocytes of the indicated *Dhrs7b* genotypes (WT and Δ/Δ) were prepared from individual tamoxifen-injected mice and divided to analyze without activation (“naive”) or after activation and culture (48 h) (“activated”). Shown are heat maps with more extensive lists of ether phospholipids and plasmalogens identified in the LC-MS-MS analyses for both genotypes in positive (A) and negative (B) ion modes. (C, D) Controlling for transgene and tamoxifen effects. Shown are LC-MS-MS results for lipids in B cells, activated or not, as indicated, of conventional non-transgenic B6 mice not injected with tamoxifen and those of tamoxifen-injected CreER^T2+^ mice (the “+/+” samples of panels A, B and of Figure 3). Bar graphs in (C) and (D) show results of these control comparisons for some of the phospholipids analyzed in prior figures panels (Fig. 3A, B).

**Supplemental Figure 4.**
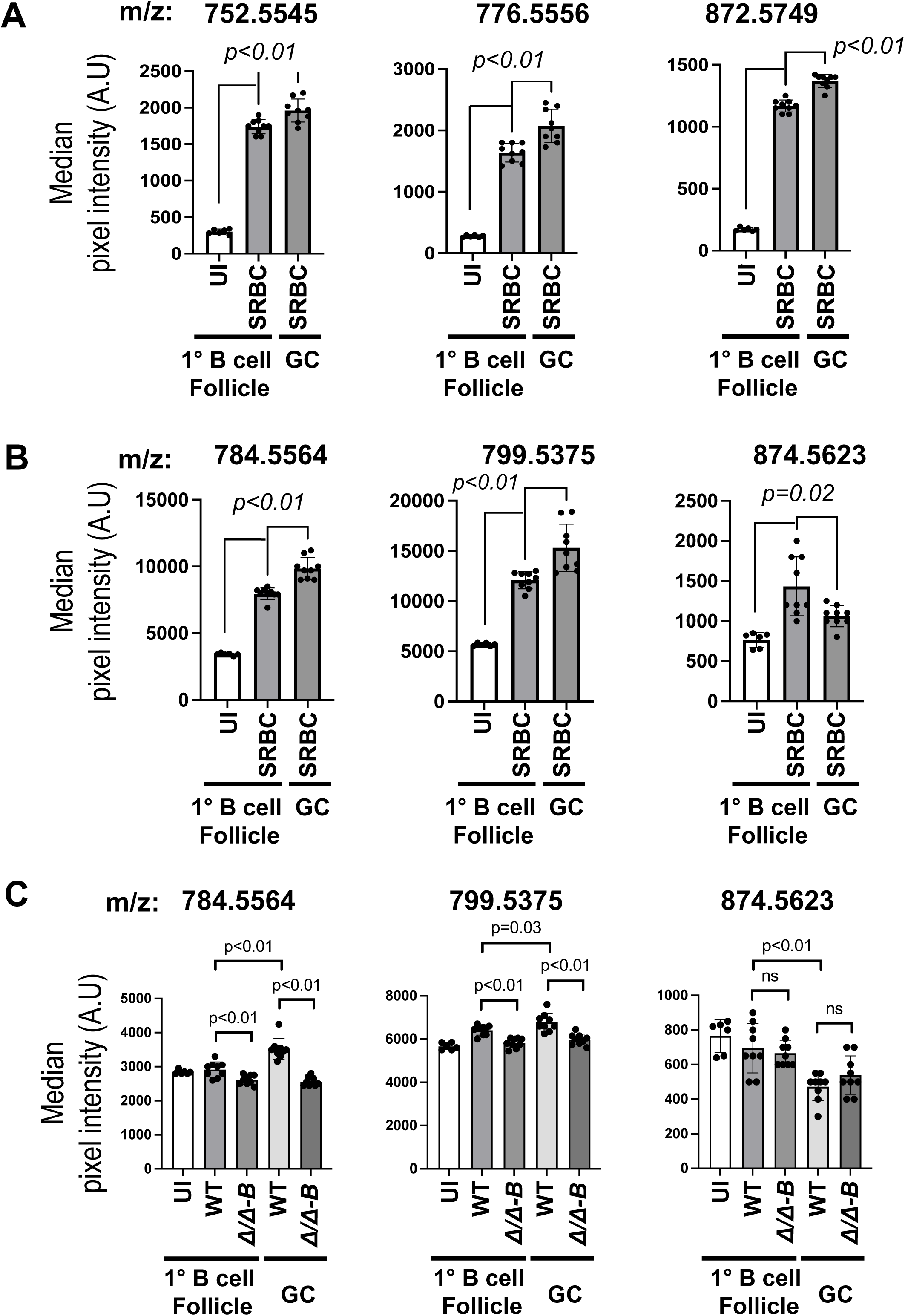
Immunization induces increases in P-lipids of both primary and secondary follicles dependent on PexRAP in B cells. (A, B) Selected representative *m/z* features Spleens of AID-GFP were harvested at 7 days after immunization with SRBC and analyzed for exact mass signatures of ether lipid species in (A) negative and (B) positive ion modes. (C) Spleens of immunized mice (tamoxifen-treated *huCD20*-CreER^T2^; *Dhrs7b^f/f^*, i.e., *Dhrs7b*^Δ/Δ-B^, and *huCD20*-CreER^T2^, i.e., “WT”, controls) were analyzed by IMS. Shown are the positive ions presented in (B) for GC vs follicles of immunized AID-GFP mice. The bar graph shows the mean (± SEM) ion intensities in lymphoid follicles and GC from WT and *Dhrs7b*^Δ/Δ-B^ spleens. Median intensity of each ion was obtained from three follicles per spleen and three GC per spleen from WT and *Dhrs7b^Δ/Δ-B^*mice (three biological replication experiments totaling seven WT and six cKO subjects). P values were calculated by Mann-Whitney U test.

**Supplemental Figure 5.**
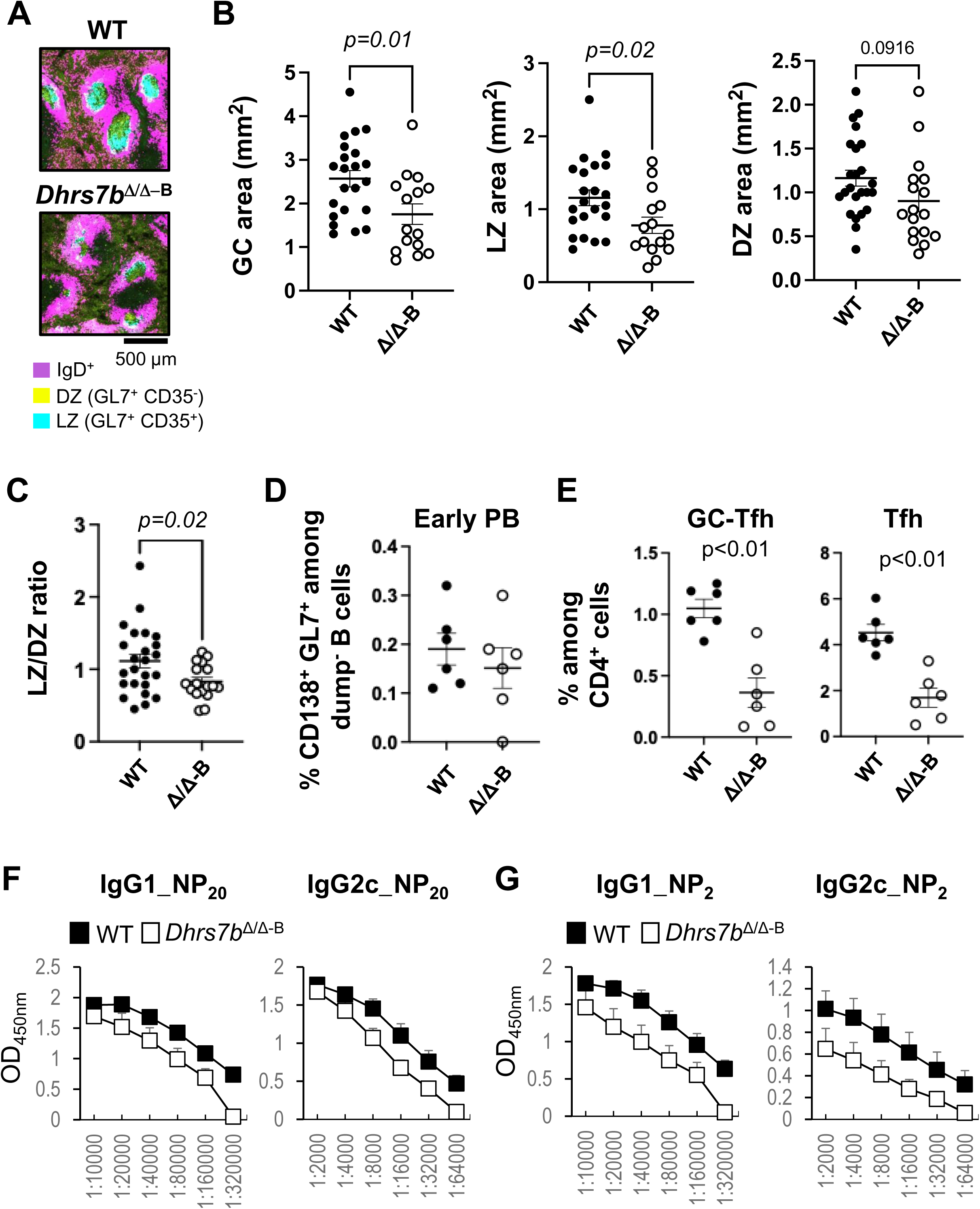
A subset of ether phospholipids in primary follicles and GC (secondary follicles) depend on PexRAP in B cells. (A) As in Fig. 4, *Dhrs7b* was inactivated by tamoxifen injections into mice bearing the huCD20-CreER^T2^ transgene, followed by immunization with SRBC and analysis. Shown are representative higher-magnification images defining the primary (IgD^+^) and secondary (IgD^lo/neg^, GL7^+^) follicles and the identification of light (CD35^+^) and dark (CD35^neg^) zones. (B-C) After immunization and harvest of the mice as shown in Fig. 4, the sizes of GC, light zones (LZ), and dark zones (DZ) in photomicrographs of sectioned and immunofluorescently stained spleens of the immunized mice (using three GC per spleen, with seven WT and five *Dhrs7b^Δ/Δ-B^*.) were quantified. (B) Spleens of immunized mice (tamoxifen-treated *huCD20*-CreER^T2^; *Dhrs7b^f/f^* and *huCD20*-CreER^T2^ controls, as indicated) were analyzed by immunofluorescence staining as in (A). Each dot represents one follicle in a given mouse. (C) Dot graph showing the ratio of LZ/DZ using the data of panel B. (D, E) Mice (*huCD20*-CreER^T2^; *Dhrs7b^+/+^, or huCD20*-CreER^T2^; *Dhrs7b*^f/f^) were treated with tamoxifen and immunized with SRBC as in Fig 4A, and were analyzed 7 d after immunization. (D) Shown are aggregated frequencies of CD138^+^ GL7^+^ early plasmablasts in B220^+^ IgD^neg^ dump^neg^ gate (dump channel = IgD, CD11b, CD11c, F4/80, Gr1, TCRb, and 7AAD). (E) The aggregated frequencies of GC-Tfh (PD-1^hi^, CXCR5^hi^) and Tfh (PD-1^int^, CXCR5^int^) among CD4^+^ T cells with the gating strategy as shown in Supplemental Fig. 1G. (x=2; 6 WT vs 6 cKO). P value was calculated by Mann-Whitney U test. (F, G) Mean (±SEM) absorbances of ELISA performed across the indicated dilutions of sera from the tamoxifen-treated mice (*huCD20*-CreER^T2^; *Dhrs7b^+/+^* or *huCD20*-CreER^T2^; *Dhrs7b^f/f^*, i.e., *Dhrs7b^Δ/Δ-B^*) of Fig. 6. Shown are the results for (F) all-(NP_20_ captured) and (G) high-(NP_2_ captured) -affinity anti-NP Ab of the IgG1 and IgG2c isotypes, as indicated.

**Supplemental Figure 6.**
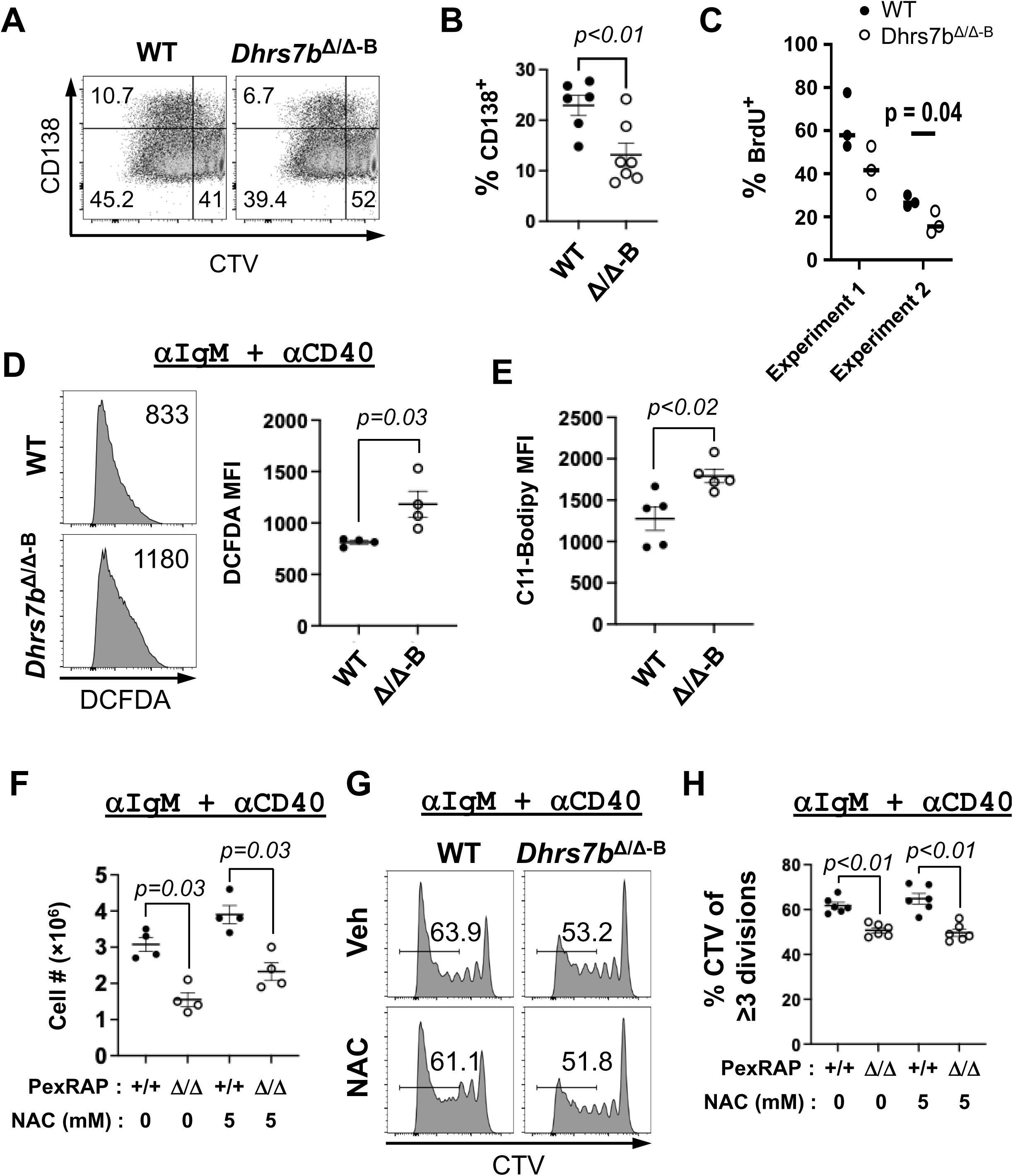
PexRAP contributes to GC B cell proliferation, ROS homeostasis and B cell population growth. (A, B) Proliferation and development of CD138^+^ progeny derived from B cells activated and cultured with anti-CD40, BAFF, IL-4, and IL-5. Shown are representative flow plots (A), and aggregated mean (±SEM) frequencies of CD138^+^ cells in viable lymphocyte from three independent experiments with 6 WT and 7 *Dhrs7b^Δ/Δ-B^* mice at t = 5dq. (C) PexRAP regulates proliferation of GC B cells. Tamoxifen-treated mice (*huCD20*-CreER^T2^; *Dhrs7b^+/+^* or *huCD20*-CreER^T2^; *Dhrs7b^ff^*, i.e., *Dhrs7b^Δ/Δ-B^*) were immunized with SRBC, and the mice were injected with BrdU as described in Methods. Shown are a dot graph aggregating all experiments’ outcomes for the frequencies of BrdU^+^ cells in GCB cells (x=2; n = 6 WT and 6 cKO). (D) WT and *Dhrs7b^Δ/Δ^* B cells were activated and cultured 3 days in anti-IgM, anti-CD40, BAFF, IL-4, IL-5, and 4-OHT. Shown are representative histogram image of H_2_DCFDA in the B cell gate (left panel), and aggregated mean (± SEM) geometric MFI of H_2_DCFDA (right panel) from two independent replicate experiment (n = 4 WT and 4 cKO). P values were calculated by Mann-Whitney U test. (E) PexRAP restrains lipid peroxidation. B cells were activated and cultured as in Fig 8E. Shown are the aggregated means (± SEM) of geometric MFI of C11-Bodipy from two independent replicate experiments (n = 5 WT and 5 cKO). (F-H) NAC mitigated the impairment of population increase for PexRAP-deficient B cells, but failed to enhance cell division. B cells of the indicated genotypes were activated with combined BCR and CD40 cross-linking as in (D), cultured 5 d in the presence or absence of NAC (5 mM), counted and analyzed by flow cytometry. (F) The dot graph shows the mean (± SEM) recovered cell number from two independent experiments. P values were calculated by Mann-Whitney U test. (G, H) Prior to activation, B cells of the each genotype were stained with CellTrace Violet (CTV). (G) Shown are flow cytometry ouputs of CTV fluorescence in the viable B cell gate for the indicated samples from one of the independent replicate pools. A measurement bar denotes the cut-off between 0-2 divisions and ≥3 divisions, with the inset numbers representing the percentage of cells that divided at least three times. (H) A dot graph displaying the individual as well as mean percentages of B cells that divided at least three times for samples of the indicated genotypes and treatment conditions. P values were calculated by Mann-Whitney U test.

**Supplemental Figure 7.**
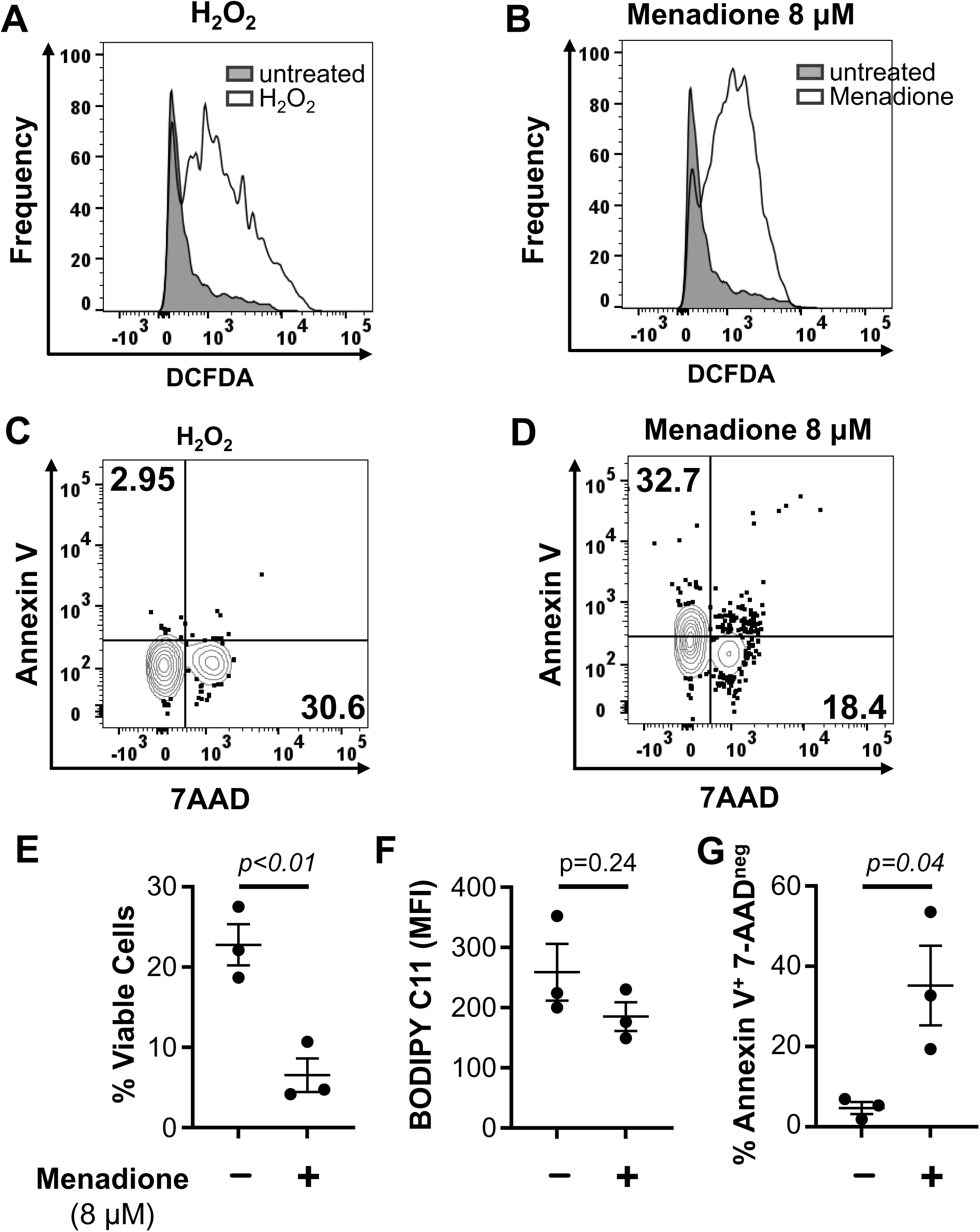
Distinct outcomes of heightened ROS elicited by H_2_O_2_ versus menadione. (A-B) Induction of ROS independent from B cell activation. Shown are representative flow plots of DCFDA in the viable B cell gate after exposure (1 h) (A) to H_2_O_2_ (200 μM) or (B) menadione (8 μM), as in (87). (C-D) Representative flow plots of annexin V vs 7-AAD for identification of early-apoptotic B cells. Cultured B cells were analyzed 3 hr after treatment with (C) 200 μM H_2_O_2_ or (D) 8 μM menadione. Inset numbers represent the fraction of B cells that are Annexin V^+^ 7-AAD^neg^ or Annexin V^neg^ 7-AAD^+^. (E-G) Shown are the results of three independent experiments to measure (E) total cell viability, (F) BODIPY C11 MFI and (G) frequencies of Annexin V^+^ 7-AAD^neg^ frequency in B cells treated (8 μM) menadione overnight (E) or for 3 h (D). P-values were calculated using the unpaired Student’s t-test.

**Supplemental Table 1. Immunization-induced increases of selected ether lipids in GC.** Shown are (a) a sample of *m/z* features characteristic of the indicated phospholipid species identified by the LIPIDMAPS database, and (b) the mean (±SEM) ion intensity / counts in IMS data generated from spleens of AID-GFP mice, immunized or not (UI), after mapping to the indicated regions of interest (primary follicles or GC, i.e. secondary follicles) using fluorescent images. Shown are ions more accumulated in GC regions compared to primary follicles (8 ions from negative ion mode, and 5 ions from positive ion mode, all p<0.05 for comparison of 1^0^ follicle to UI (c), or GC to 1^0^ follicle (d). P values were calculated by Mann-Whitney U test.

**Supplemental Table 2. Impact of PexRAP on relative quantities of selected lipid species in primary and secondary follicles (GC).** As in Table 1 except that B cells of immunized mice were either WT or lacked PexRAP (cKO, i.e., *Dhrs7b*^D/D-B^). Shown are (a) a sample of *m/z* features characteristic of the indicated phospholipid species. (b) the mean ±SEM ion intensity counts in IMS data after mapping to the indicated regions of interest as in Table 1. (c-f) indicate that p<0.05 for the null hypothesis in considering a difference between WT mouse spleens after immunization versus UI controls (c), 1^0^ B cell follicles in spleens of immunized WT vs *Dhrs7b*^Δ/Δ-B^ mice (d), GC vs in 1^0^ B cell follicles in spleens of immunized WT mice (e), and GC of immunized WT vs *Dhrs7b*^Δ/Δ-B^ mice (f) in the ions counts for designated *m/z* features. P values were calculated by Mann-Whitney U test.

